# Affinity-selected peptide ligands specifically bind i-motif DNA and modulate c-Myc gene expression

**DOI:** 10.1101/2025.05.30.656635

**Authors:** Dilek Guneri, Summer Rosonovski, Effrosyni Alexandrou, Shuang Chen, Jessica King, Christopher A. Waudby, Shozeb Haider, Christoper J. Morris, Zoë A. E. Waller

## Abstract

c-Myc is an oncogene that is dysregulated in ~70% of cancers. Its multifaceted function complicates effective drug targeting of the protein. i-Motif DNA structures in gene promotor regions have gained attention for their potential role in modulation of gene expression. These include the i-motif formed by the cytosine-rich sequence that lies upstream of the key P1 promotor of the c-Myc gene. Currently, selective ligands interacting with i-motif structures are limited. Here, peptide ligands for the i-motif from the promoter of c-Myc were identified *via* phage display. Hit peptides were filtered for selective binding to i-motif structures over other DNA structures using displacement assays and DNA melting experiments. Two lead peptides were found to produce dose-dependent changes on c-Myc gene expression after delivery into HEK293 cells expressing a c-Myc luciferase reporter construct. These leads may be used as chemical tools for the manipulation of c-Myc i-motif *in vitro* and have potential to be developed into cell-permeable peptidomimetics for delivery *in vivo*.

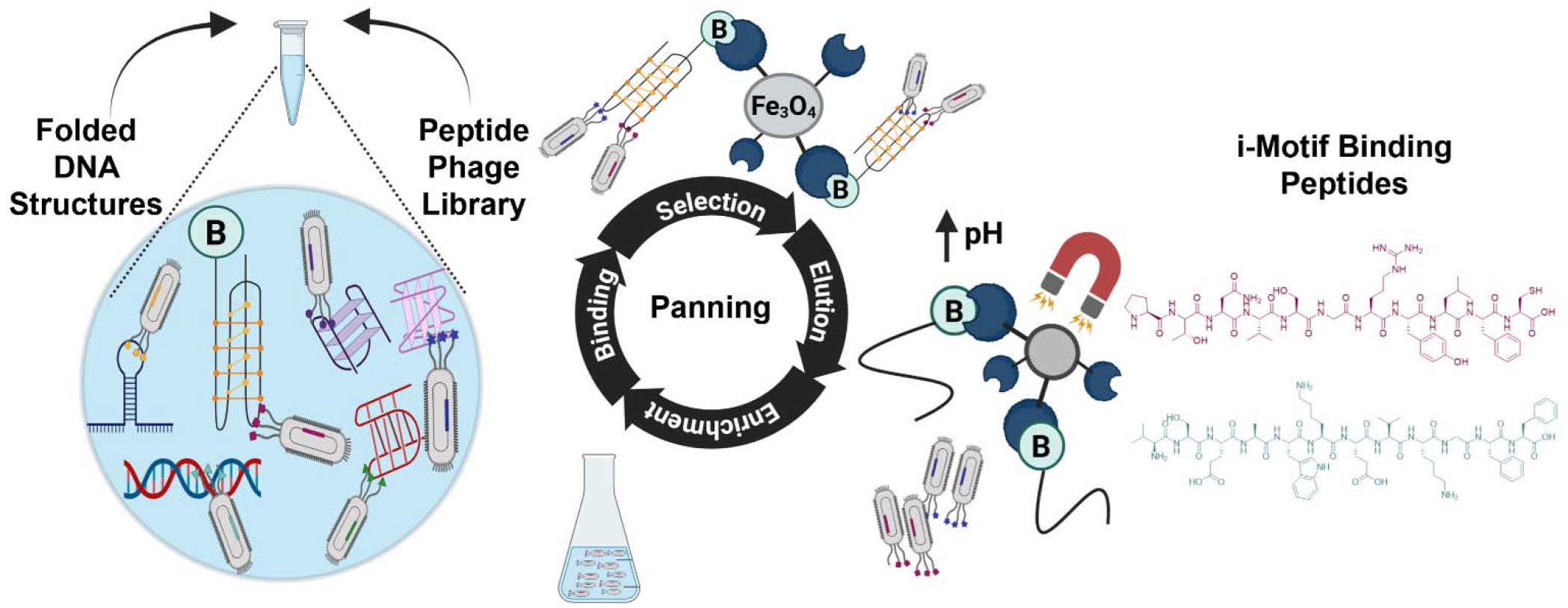

## Introduction

The proto-oncogene, c-Myc, is a multifunctional transcription factor which controls various downstream genes and coordinates a range of cellular processes in both health and disease.^1^ Elevated expression of c-Myc in cancer occurs through several mechanisms (e.g. gene amplification, chromosomal translocation) and results in elevated tumour aggression and poorer clinical outcomes.^2^ Substantial experimental evidence has established that reducing overactive c-Myc can lead to cancer regression.^1, 3-5^ One approach to direct targeting of the Myc protein is achieved by interference of the Myc-Max heterodimerisation. The peptide/protein candidates, OMO-103 and IDP-121, have progressed to Phase I clinical testing. OMO-103 dose-escalation trial results indicate that this is a tractable approach that can deliver anti-tumour activity across different tumor types.^6^ A significant challenge for all protein/peptide-based drugs is the requirement to deliver an intracellular dose that is sufficient to neutralize all copies of the c-Myc protein in a cancer cell. These have been estimated to number 30,000 copies per cell.^7^ Reports of siRNA-mediated knockdown of c-Myc mRNA levels indicate that this approach has merit, yet efficient intracellular delivery of siRNA can be similarly challenging and inefficient. A contrasting and arguably more potent approach involves targeting the c-Myc gene directly such that MYC levels are suppressed pre-transcriptionally through blockade of key protein-DNA promoter interactions. This approach is particularly appropriate considering the short half-life of both c-Myc mRNA (10 minutes) and protein (20 minutes).^8^

c-Myc transcription is largely controlled by the P1 and P2 promoter sites.^9^ Upstream of the P1 promoter regulatory region there is a nuclease hypersensitivity element (NHE)-III_1_ that is prerequisite for approximately 90% of c-Myc gene transcription.^10-12^ This element contains GC-rich sequences capable of forming non-canonical DNA structures: i-motifs (iMs) and G-quadruplexes (G4s). iMs are formed from cytosine-rich sequences, zipped together through hemi-protonated cytosine-cytosine base pairs^13, 14^ whereas G4s on the opposing strand form in guanine-rich sequences, stabilized by Hoogsteen hydrogen bonding and physiologically-relevant cations.^15^ G4-formation within the NHE III_1_ region has been extensively studied, showing that destabilizing G4 structure leads to increased c-Myc transcription and stabilizing the G4 repressed transcription.^16-18^ c-Myc transcription is proposed to be initiated by the negative supercoiling of the NHE-III_1_ region that facilitates the G4 formation and creates a transient single-stranded region recognizable for elements of the transcription machinery. This may include helicases able to resolve the G4 structure and promote c-Myc expression.^19^ However, negative supercoiling alone may not be the key mechanism to facilitate G4 formation.^20^ Further insights into the regulatory region of the c-Myc promotor region are required to facilitate the development of novel ligands and potential drugs. This includes exploration and study of the potential of iM-formation on the opposing strand.^21^ iM-forming sequences are well-characterized to be able to form *in vitro* at neutral pH^22, 23^ and have been shown to be present and involved in transcription and proliferation, in cells.^24-26^ More recently, there have been reports of iM-binding small molecules,^27^ but selectivity remains an issue for iM ligands. Indeed, even many G4-binding ligands have also been found to interact with diverse iM structures.^28, 29^ Consequently, the discovery of specific iM-binding ligands is crucial to develop both target-specific chemical tools and future therapeutics both for c-Myc and other iM-forming regions in the genome.

Specific nucleoproteins have been found to bind both iMs or G4 structures and subsequently modulate gene expression levels. Heterogeneous nuclear ribonucleoprotein K (hnRNP-K) is an RNA/DNA binding protein that binds to single-stranded, cytosine-rich sequences. It was shown by Hurley and co-workers to bind and unfold the iM-forming sequence of c-Myc and to increase c-Myc transcription.^30^ This was indirectly validated by others who observed that hnRNP-K gene-knockdown reduced MYC protein levels.^31^ More recent work has demonstrated different ways to target specific DNA G-quadruplexes, including c-Myc, using CRISPR-guided G-quadruplex-binding proteins and ligands.^32, 33^ Drawing inspiration from the binding activity of nucleoproteins, we aimed to identify peptidic ligands that could regulate quadruplex biology. Unlike non-selective small molecules and large antibody species, these peptide ligands would occupy a molecular size niche with potential for increased specificity compared to small molecules and improved intracellular delivery capabilities compared to antibodies. Here, we deployed phage display to select peptide ligands that bind selectively to the iM-structure in the c-Myc promotor.

## Materials and Methods

### Chemicals

were purchased from Sigma Aldrich/Merck, were of analytical or molecular biology grade and used without further purification.

### Oligonucleotides

(listed in Tables S1, S2) were purchased from Eurogentec and were RP-HPLC purified. Solid DNA samples were initially dissolved in ultrapure water to a concentration of 1 mM. Stock concentrations were confirmed using the extinction coefficients provided by Eurogentec and their UV absorbance at 260 nm is determined with a Nanodrop ND-1000 spectrophotometer. The samples were prepared as 10 µM DNA in 10 mM sodium cacodylate (NaCaco) buffer at the specified pH and thermally denatured at 95□°C for 5□minutes and allowed to anneal slowly to room temperature overnight. pH 6.6 was used for most of the biophysical studies as it is the transitional pH of c-Myc, the pH where 50% of the sequence is folded into iM.^30^

### Phage display of peptides and c-Myc i-motif ligand selection

The NEB Ph.D.^TM^-12 library, containing *ca*. 10^9^ randomized, linear peptide sequences was panned against c-MycC52 i motif following the protocol described in the NEB Phage Display instruction manual. Key buffers used in the library panning were: Incubation buffer (PBS pH 6.0), wash buffer (PBS pH 6.0 with 0.01-0.05% Tween 20), blocking buffer (PBS pH 6.0 with 5 mg/mL BSA), and elution buffer (PBS, pH 7.4) (Table S3). Phage display library panning involved solution-phase panning with a 5′-biotinylated c-MycC52 target and target capture on High-Capacity Streptavidin coated plates. PBS was the DNA buffer of choice for phage display as has a good buffering range across pH 5.8-7.4. c-MycC52 is unfolded at pH 7.4, fully folded at pH 5.8, and 50% folded at pH 6.44 (Figure S1), consistent with previous reports by Hurley and co-workers.^30^ PBS pH 6.0 was chosen as the incubation buffer as this is slightly more acidic than the transitional pH of c-MycC52 in PBS (Figure S1). Table S3 summarises the three rounds of phage display screening that were undertaken. Target c-MycC52 oligonucleotide was used alone in round one. Unbiotinylated competitor oligonucleotides were used in rounds two and three at the concentrations indicated in Table S3. At the end of each screening round phages were enumerated by titration with E. coli ER2738 and amplified in E. coli ER2738 as per NEB’s phage display manual. At the end of round three, twenty-five individual phage plaques were harvested from IPTG/X-Gal plates and amplified on a 5 ml scale for 4 hours. ssDNA was isolated after iodide denaturation and ethanol precipitation. DNA sequenced using Sanger sequencing (Mix2Seq Eurofins) using the −96 gIII sequencing primer provided by NEB.

### Peptides

were purchased from Cellmano Biotech (Hefei, China) as RP-HPLC purified (> 95%) with N-terminal acetylation and C-terminal amidation to mimic the phage-displayed fusion peptides. Stock solutions of the peptides were made at 5 mg/mL in ultrapure water or DMSO (if water insoluble) and were stored at −20□°C. Subsequent dilutions were made in the appropriate assay buffer. Stock concentrations were confirmed using the peptide extinction coefficients and UV absorbance on Agilent Technologies Cary 4000 UV-VIS spectrometer. The Nick Anthis’ Protein Parameter Calculator was used to find the extinction coefficient either at 280 nm if the sequence contained any tryptophan or tyrosine residues or at 205 nm if the sequence contained neither of these amino acids.^34^

### Fluorescent Indicator Displacement (FID) experiments

used in this publication are based on the first FID assay developed for i motif DNA.^35^ The FID experiments were performed on a BMG CLARIOstar plate reader using 96-well, solid black flat bottom plates. Thiazole Orange (TO) was the indicator, and the stock solution was prepared at 10 mM in DMSO. The TO stock was diluted into the appropriate buffer to 2 µM. Each well had 9 µL of the 2 µM TO solution added and excited at 430 nm with fluorescence emission at 450 nm measured; this was normalized to 0% to account for background fluorescence. 1 µL of 90 µM DNA was added to each well and shaken in the plate reader for 30 seconds using double orbital shaking at 700 rpm and left for 10 minutes to equilibrate. Following equilibration, fluorescence emission was measured and normalized to 100% representing maximum fluorescence. Peptides were titrated in 25 µM in 0.9 µL additions until 200 µM into each well in triplicate until 200 µM of peptide was reached, and fluorescence emission was measured following each addition. Fluorescence emission for each well was normalized between 0 and 100% which was taken away from 100 to give the percentage of displacement. The data was analyzed in Origin data analysis software and fitted with a hyperbolic dose-response curve from which the DC_50_ value was interpolated.

### Circular Dichroism (CD) melting experiments

were performed in Jasco J-1500 spectropolarimeter with 10 µM DNA samples in 10 mM sodium cacodylate buffer at pH 6.6 using a 1 mm path length quartz cuvette. Initially two repeats of CD melting full spectrum ranges were taken (from 230 nm to 320 nm) for c MycC27, c-MycC52, c-MycG27, c-MycG52, and dsDNA (Table S1) measuring the unfolding of the DNA structures from 5°C - 95°C in presence of 10 molar equivalent DMSO, Pep-PTN or Pep-VSE as well as Pep-SLC and buffer for dsDNA. The samples were kept at 5°C for 5 minutes before starting the melt with 1°C/min increase in temperature, 0.5°C data interval and 60 seconds holding time at each target temperature. Four scans were accumulated data pitch at 0.5 nm, scanning speed of 200 nm/min, 1 second response time, 2 nm bandwidth, and 200 mdeg sensitivity. Furthermore, two CD melting experiment repeats were performed recording C-rich c-Myc at 288 nm and 320 nm, G-rich c-Myc at 264 nm and 320 nm, and dsDNA at 253 nm and 320 nm with the settings as described for full spectrum measurements. Data was zero corrected to 320 nm and baseline drift. The melting temperature (*T*_M_) was concluded using the Boltzmann or biphasic fitting curve on the folded fraction data using GraphPad Prism version 10.1.2. Data was processed as Mean ± SEM (n=4) and One-way ANOVA followed by Bonferroni post-hoc test to determine significant changes between peptides and controls.

### UV titration

were recorded using an Agilent Cary-60 UV/VIS spectrometer with a 10 mm path length black-walled quartz cuvette. Oligonucleotides (c-MycC27, c-MycC52, c-MycG27, c-MycG52, DAPc, DAPg, and dsDNA; sequences listed in Table S1) were annealed as 250 µM stock solutions in 10 mM sodium cacodylate buffer (pH 6.6), in the presence or absence of 100 mM KCl. Spectra were recorded at room temperature over a wavelength range of 300–500 nm with 1 nm intervals. Buffer-only spectra were recorded and subtracted from each dataset as blanks. Peptide ligands were prepared as 1 mM stock solutions in DMSO and added to the cuvette at a final concentration of 10 µM in buffer prior to DNA addition. DNA stock was diluted to 50 µM in buffer and was added stepwise to the peptide solution to reach 2.5 µM final concentration, followed by additions from the 250 µM DNA stock to reach final DNA concentrations of 30 µM or 50 µM. The absorbance at 350 nm was monitored and plotted against DNA concentration. Binding curves were fitted using a one-site binding model in GraphPad Prism version 10.1.2, and K_d_ values were determined from normalized data to represent the DNA-ligand bound fraction.

### NMR

1H NMR spectra of c-MycC52 were recorded at 283 K using a Bruker Avance NEO 600 MHz spectrometer equipped with QCI-F cryoprobe operating Topspin 4.5.0. Spectra were recorded with excitation sculpting, at 27.75 ppm spectral width, 1 s acquisition time and 1 s recycle delay. Spectra of the imino proton region were acquired with a 1D SOFAST excitation scheme with a 1293 μs Pc9_4_120 excitation pulse and 720 μs Reburp refocussing pulse, centered at 13 ppm, a 29.75 ppm spectral width, 115 ms acquisition time and 100 ms recycle delay. Samples were annealed at a final concentration of 50 µM in 10 mM sodium cacodylate (pH 6.6). 10% D2O and 0.01% DSS was added immediately before recording NMR spectra experiments. Spectra were processed with exponential line broadening and referenced to the DSS chemical shift. The chemical shift of the cacodylate buffer resonance was used to monitor potential pH changes between samples.

### Cell Studies

HEK293 cells (ATCC CRL-1573™, passages 20–28), MCF7 cells (ATCC HTB-22™, passages 21–28) and Panc-1 cells (ATCC CRL-1469™, passages 15–22) were cultured in DMEM medium supplemented with 10% fetal bovine serum (FBS; Gibco, UK) and maintained under standard cell culture conditions (37°C, 5% CO□). Cells were passaged every 3–4 days at ~80% confluency. For experiments, the cells were into 6-well plates at a density of 1 × 10□ cells/well in DMEM containing 10% FBS. At ~80% confluency, HEK293 wild-type cells were co-transfected with the c-Myc promoter (Del-4) Firefly luciferase plasmid (a gift from Bert Vogelstein; Addgene plasmid #16604; http://n2t.net/addgene:16604; RRID:Addgene_16604) and a control Renilla luciferase reporter vector (Promega, pRL-TK Vector; GenBank® Accession Number AF025846) at a 3:1 plasmid ratio. In parallel, the pGL410_INS421 plasmid (a gift from Kevin Ferreri; Addgene plasmid #49057; http://n2t.net/addgene:49057; RRID:Addgene_49057), which contains the human insulin promoter with 2.5 insulin-linked polymorphic region (ILPR) repeats regulating firefly luciferase expression,^36^ was transfected as a control for promoter activation. Plasmid DNA and Lipofectamine™ 2000 (Invitrogen, UK) were mixed at a 1:1 ratio and incubated at room temperature for 15 minutes before transfecting HEK293 cells. The following day, HEK293 transfected with reporter vectors for c-Myc or ILPR were re-seeded into white and transparent 96-well plates (Greiner Bio-One, Germany) at a density of 5 × 10□ cells/well in DMEM containing 2% FBS. Cells were treated in duplicates with 1 µM, 5 µM, or 10 µM of Pep-PTN or Pep-VSE, or with 0.2% DMSO as a vehicle control, in the presence or absence of Nanocin PRO (Tecrea, UK) as a peptide delivery reagent. In addition, HEK293 wild-type, MCF7 cells and Panc-1 cells were also seeded at 5 × 10□ cells/well into transparent 96-well plates in DMEM with 2% FBS for cell proliferation assays. Cell proliferation was assessed at 4 h and 24 h post-treatment using the CellTiter 96® Aqueous One Solution Cell Proliferation Assay (Promega, USA). For this, 20 µL of reagent was added per well, and absorbance at 490 nm was measured after 4 h and 24 h using a SpectraMax iD3 plate reader (Molecular Devices, UK). Blank values (media without cells) were subtracted, and readings were normalized to DMSO-treated controls. In parallel, Dual-Luciferase Reporter Assays (Promega, USA) were performed in white 96-well plates to determine promotor activation regulated by c-Myc or ILPR promotor regions. Firefly luciferase signals were normalized to Renilla luciferase signals and further normalized to the 0.2% DMSO control condition. All data represent the mean ± standard deviation (SD) from six to eight biological replicates. Statistical significance was determined using one-way ANOVA followed by Holm Šídák post-hoc analysis performed in GraphPad Prism version 10.1.2.

### Computational Studies

A model of the c-Myc i-motif with a sequence of 41 nucleotides was built (5′-CTTCTCCCCACCTTCCCCACCCTCCCCACCCTCCCCATAAG-3′). This model was based on the intramolecular iM structure from the insulin-linked polymorphic region^36^ and folding conformations proposed by Sutherland *et. al*.^30^ (Fig. S21). Adaptive bandit enhanced sampling molecular dynamics simulations^36, 37^ were employed to sample the loop and flank conformations of this model. Markov state models were then built to cluster metastable states using the PyEMMA package.^38^ From these, the two most stable states were chosen to carry out molecular docking with the PTN and VSE peptides. De novo 3D structure predictions for these peptides were performed, using the PEP-FOLD3 web server (https://bioserv.rpbs.univ-paris-diderot.fr/services/PEP-FOLD3/). Docking was subsequently performed using the HDOCK web server (http://hdock.phys.hust.edu.cn/). Full details of all the protocols are listed in the Supplementary Information.

## Results and Discussion

Phage display is a recombinant screening technique that permits the rapid screening of libraries with diversity up to 10^9^ unique ligand peptides or antibody fragments against a given target such as iM and G4 structures.^24, 39^ We screened a linear peptide library (NEB Ph.D.TM12) against the biotinylated 52 bp C-rich c-Myc promoter sequence (c-MycC52, 5′-CTT-CTC-CCC-ACC-TTC-CCC-ACC-CTC-CCC-ACC-CTC-CCC-ATA-AGC-GCC-CCT-CCC-G-3′). This longer-length c-Myc sequence has been shown to be necessary for effective transcriptional firing from the P1 and P2 promoters.^30, 40^ The selection pressure for each round of panning was increased by reducing the amount of biotinylated target c-MycC52 used and adding non-biotinylated competitor DNA sequences. Round one of panning used 10 pmol of c-MycC52 alone, without competitors. Rounds two and three reduced the amount of c-MycC52 target and then added non-biotinylated competitors. Competitors included in round two included three different types of G-quadruplexes, a short double stranded sequence, Holliday junction, single-stranded RNA, a C-rich hairpin which does not form iM and calf-thymus DNA. Round three had a range of competing iM-forming sequences which represent a range of C-stack and loop lengths, with varying pH stabilities: sequences from the human telomere, the ILPR and the promoter regions of ATXN2L, DAP and Hif1α (Table S2,S3). DNA Sequencing was performed at the end of round three to determine the sequence of the selected peptides, which revealed in total five different peptides (Table 1). Full sequences, structures and purity of the peptides purchased and subsequently used are provided in Table S4.

**Table 1.** Peptide sequences determined from three rounds of panning of Phage Display against c MycC52 using the NEB Ph.D.TM12 linear randomized library. Conserved sequence shown in bold; consensus sequence underlined.

| Peptide Ligand | Amino acid sequence |
| --- | --- |
| Pep-EIE | EIEY**TDHMK**ELG |
| Pep-PTN | PTNVSGRNYLFC |
| Pep-RVS | RV**STDHMK**GRGG |
| Pep-SLC | SLCDIIRIEKVR |
| Pep-VSE | **VSE**AWKEVKGFF |

The most enriched peptide was Pep-EIE, which shared a TDHMK motif with Pep-RVS. Three peptides contained the consensus sequence: (S/T)(D/E)XX^2^X^3^ where X is any amino acid, X^2^ is a large amino acid and X^3^ is a basic amino acid. Pep-SLC included the closely related sequence, DIIR. Pep-PTN and Pep-SLC both included at least one arginine residue, which are typically depleted in the particular phage display library used.^40^ Conventionally, arginine residues favor binding to pyrimidine-rich DNA regions, as evidenced by the abrogated binding of NM23-H2 to single stranded DNA after the substitution of a key arginine for alanine, reducing interactions with the phosphodiester backbone.^41^

The relative binding of the five peptide ligands to the folded c-MycC52 iM were assessed via a fluorescent indicator displacement (FID) assay^30^ using thiazole orange (Fig. 1). These experiments were performed in 10 mM sodium cacodylate at pH 6.6, the transitional pH of the c-Myc iM-forming sequence. As iM-formation is reduced at higher pH and higher ionic strength,^42^ additional stabilizing cations were avoided to allow the highest proportion of iM formation at the highest pH, to represent as closer to physiological pH conditions as possible.

**Figure 1.**
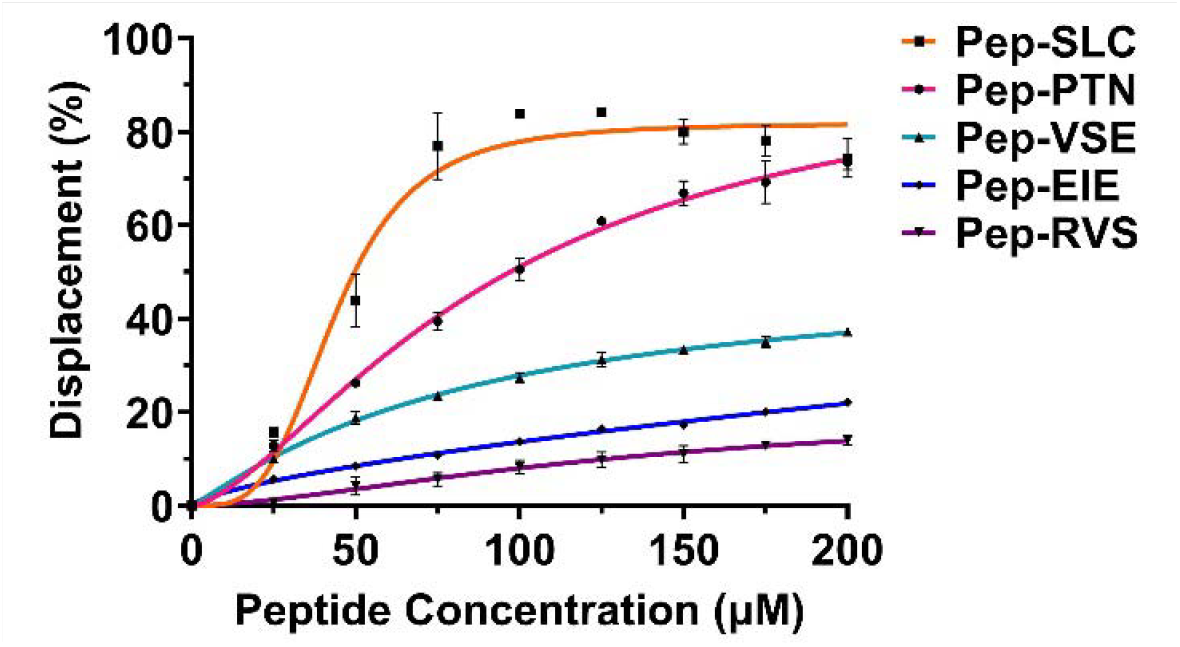
Concentration-dependent interaction of peptide with c-MycC52 i-motif determined via FID assay. The assay was performed with 1 µM c-MycC52 in 10 mM Sodium Cacodylate, pH 6.6, in the presence of 2 µM thiazole orange. Peptide ligands were added in increments of 25 µM up to a final concentration of 200 µM. Data are presented as Mean ± SD (n=3) and fitted using a Hill-Slope dose-response curve.

All five peptides demonstrated concentration-dependent intercalator displacement from the c-MycC52 iM. The relative binding affinity was ranked as SLC>PTN>VSE>EIE>RVS. All were able to show displacement of TO from the c-MycC52 sequence, with average displacement values at 100 µM spanning 5-85%. It is important to note that the thiazole orange assay gives an indication of displacement, rather than direct binding. Accepting that this method only determines which peptides displace the TO, and that the peptides may bind also other sites on this extended iM, we took the top three peptides SLC, PTN and VSE forwards for further study at this stage.

To determine the effects of peptide binding on iM stability we measured DNA melting points (T_M_) using circular dichroism (CD)-based melting experiments. The three remaining peptides were initially tested for off-target binding to double stranded (ds) DNA. Pep-SLC destabilized dsDNA by 5°C (Fig. S2A, Fig. S3). Pep-PTN and Pep-VSE had no effect on dsDNA melting compared to the solvent controls (Fig. S2B, Fig. S4). As we were interested in characterising peptides with the maximum specificity, we elected to exclude Pep-SLC from further characterisation at this time, though this peptide may still have utility for targeting c-Myc. We tested the thermal stability of c-MycC52 iM and a shorter analogue c-MycC27 (5′-CCT-TCC-CCA-CCC-TCC-CCA-CCC-TCC-CCA-3′) (Fig. S5) as well as their complementary G4-forming sequences (c-MycG52 5′-CGG-GAG-GGG-CGC-TTA-TGG-GGA-GGG-TGG-GGA-GGG-TGG-GGA-AGG-TGG-GGA-GAA-G-3′) and c-MycG27 (5′-TGG-GGA-GGG-TGG-GGA-GGG-TGG-GGA-AGG3′) respectively in Fig. S6, in the presence of 10 molar equivalents of Pep-PTN or Pep-VSE. The stabilisation temperatures are summarised in Table S5 and full meting curves are shown in Fig. S2-4 and S7-10. Both c-MycC27 and c-MycC52 sequences exhibited a double transition in the CD melts, implying two folded populations with different stabilities (Fig. S5). This was not unexpected, as both sequences have the capabilities of folding into different structures because of their length.^39^ We determined the *T*_M_s of both species due to the likely existence of both iM topologies *in vivo*. The c-MycC27 and c-MycG27 sequences were neither stabilised nor destabilised by Pep-PTN nor PEP-VSE (Table 2, and Fig. S5B & Fig S7B). In contrast, the peptides significantly modulated the thermal stability of the biologically relevant c-MycC52 and the complementary G-rich sequence (p<0.001***). Specifically, both iM populations formed by c-MycC52 showed significant destabilisation in presence of Pep-VSE (Table 2, Figs. S5A). Interestingly, the *T*_M_ decrease of the c-MycC52 sequence was greater for the more stable iM population, with a Δ*T*_M_ of −14±0.5°C (p<0.001***, see Fig S5A).

**Table 2.** Destabilisation and binding properties of Pep-PTN and Pep-VSE. CD melting analysis of 10 µM c-MycC27 and c-MycC52 and their complimentary c-MycG sequences in 10 mM Sodium Cacodylate at pH 6.6 buffer in the presence of 10 molar equivalents of Pep-PTN, Pep-VSE or the DMSO solvent control. For c-MycC52, where there were two transitions, the top Δ*T*_M_ indicates the one observed at lower temperatures and the bottom value is the second transition, at higher temperature. All data are presented as Mean change in the higher Tm ± SEM (n=4). Significance tested with a one-way ANOVA with Bonferroni post-hoc analysis. * p<0.05, ** p<0.01, *** p<0.001, highlighted in bold. Binding constants determined by UV-titrations in 10 mM sodium cacodylate at pH 6.6. Values represent the average of three experiments ± standard deviation. A *K*_d_ >50 indicates cases where evidence of binding was observed but did not plateau at the concentration range of the experiment. NS = non-specific binding indicates where no significant pattern of binding was observed at the concentration range of the experiment.

| Peptide | $\Delta T_M (^\circ\text{C})$ | | | | | $K_d (\mu\text{M})$ | | | | |
| --- | --- | --- | --- | --- | --- | --- | --- | --- | --- | --- |
|  | c-MycC27 | c-MycC52 | c-MycG27 | c-MycG52 | DS | c-MycC52 | c-MycG52 | DAPc | DAPg | dsDNA |
| PTN | $0 \pm 0.5$<br>$0 \pm 0.5$ | $-1 \pm 0.5$<br>**$-2 \pm 0.3^*$** | $-1 \pm 0.5$ | **$+2 \pm 0.3^*$** | $0 \pm 0.6$ | $0.93 \pm 0.08$ | $> 50$ | NS | NS | NS |
| VSE | $0 \pm 0.6$<br>$-1 \pm 0.6$ | **$-3 \pm 0.2^{**}$**<br>**$-14 \pm 0.5^{***}$** | $-1 \pm 0.5$ | **$+4 \pm 0.1^{**}$** | $0 \pm 0.8$ | $0.43 \pm 0.03$ | $> 50$ | NS | NS | NS |

In contrast, Pep-PTN caused a modest, 2±0.3°C decrease in *T*_M_ (p<0.05) of the more stable iM population with no significant change to lower *T*_M_ value. The complementary c-MycG52 sequence was modestly stabilised by both Pep-PTN (+2°C, p<0.05) and Pep-VSE (+4°C, p<0.01**) (Table 2 & Fig. S6A). To the best of our knowledge these are the first examples of ligands to target the long c-MycC sequence. There are a few ligands that have been found to target shorter versions of the sequence. For example, Li *et. al*. developed compound B19 which was found to stabilise the shorter c-MycC27 sequence (Δ*T*_M_ 11.5 °C),^43^ Saha *et. al*. found a compound that stabilised a 33-mer variant of c-Myc (Δ*T*_M_ 29.7 °C)^44^ and Debnath *et. al*. found a compound that stabilised the same sequence with a Δ*T*_M_ value of 32 °C.^45^ Porzio *et. al*.^*46*^ examined compounds against a 22-mer version and their best compound against that sequence of c-Myc showed a Δ*T*_M_ of −27.5 °C.

To support the experiments on the stabilising properties of the peptides, we also performed direct binding measurements of Pep-VSE and Pep-PTN using UV titrations. Binding curves are provided in Fig.S11 and the *K*_d_s are in Table 2. These experiments reveal tight binding of both PTN (0.93 ± 0.08 µM) and VSE (0.43 ± 0.03 µM) compared to other DNA structures. Although binding was detected for both peptides against c-MycG52, the G-quadruplex forming sequence in the complementary strand, there was not a plateau at the concentrations examined, so we have estimated a lower limit of 50 µM as the potential dissociation constant. Binding measurement against the iM- and G4-forming sequence from the promoter region of death associated protein (DAPc and DAPg respectively) and double-stranded DNA showed no concentration-dependent or saturable binding event to either peptides, indicating non-specific binding under tested conditions. As much of the work we performed with c-MycC52 was done in the absence of additional stabilising monovalent cations, to enable experiments to be performed at the highest reasonable pH possible, we also determined the binding in the presence of 100 mM KCl, and these binding affinities were also within the same ranges (Table S6 and Figures S11-S13). This gives about 50-fold selectivity (Pep-PTN) and 100-fold selectivity (Pep-VSE) for c-MycC52 against the complementary G-quadruplex forming sequence, or indeed any other DNA structure examined. Despite the demonstrable importance of the extended five-tract C-rich sequence within the c-Myc promoter,^30^ most studies looking at ligand binding use the shorter sequences. Some examples include Dash and co-workers, who demonstrated ligands PBP1 and PBP2 that could bind the short c-Myc sequence with *K*_d_s of 2.4 and 9.5 µM respectively^45^ and that ligand 3be has an apparent *K*_d_ value of 0.25 μM.^44^ Mitoxantrone and analogues showed a range of *K*_d_ values for these ligands against the short c-Myc iM forming sequence of between 6.8 and 38 µM,^47^ tobramycin was also found to bind c-Myc with a *K*_d_ of 13 μM.^35^ Analogues of bisacridines have reported binding affinities of between 1 and 26 μM^48^ and acridone derivatives show binding between 4.6 and 6.8 μM.^49^ The *K*_d_ for these peptides against c-MycC52 (PTN = 0.93 ± 0.08 µM and VSE = 0.43 ± 0.03 µM) is less than all but one of these examples of ligands binding c-Myc. The iM-specific antibody has un-matched affinity for the c-Myc iM structure (*K*_d_ = 409 pM), but data are reported only for binding to the shorter c-Myc sequence. Although the iM-targeting antibody is specific for iM as a structure, it also bind multiple types of iM-structures.^24, 50^ Here these peptides demonstrate unprecedented specificity for the iM from c-Myc under the examined conditions.

The profound iM-interacting capabilities of these peptides prompted us to examine if they could modulate gene expression in cells. The widely used Del-4 c-Myc promoter reporter construct was selected to investigate the biological effects of the peptide delivery into cells. The large molecular mass (>1500 g/mole) of the peptides at physiological pH prompted us to use Nanocin Pro^®^ to enhance cellular uptake. Relative c-Myc promoter activity was measured in a dual reporter gene assay by normalising Firefly expression, which is proportional to c-Myc expression, and *Renilla* luciferase expression as an internal control (Fig. 2A). Treatment with both Pep-PTN and Pep-VSE caused a concentration-dependent decrease in luciferase activity up to a 16-36% increase across a 1-10 µM range (p<0.001 - 0.01) compared to the vehicle only control (Nanocin Pro^®^ + 0.2% DMSO). This indicated that the peptides cause a decrease in c-Myc promoter activity. This is in-line with the hypothesis proposed by Hurley and co-workers in their study of the C-rich sequences from the promoter region of Bcl-2.^51^ In their study they showed that stabilisation of iM structure resulted in an increase in expression of Bcl-2. Our peptides have the opposite biophysical properties, and the opposite biological effects. Experiments were also performed without Nanocin Pro^®^ delivery agent and also showed that the peptides do still have an effect, although reduced compared to the treatments with Nanocin Pro^®^, in the absence of this type of delivery agent (Fig.S14). We were interested in understanding the specificity of the effects on c-Myc promoter activity, so also examined the effects of the peptides against an analogous reporter gene from the insulin promoter, which contains the iM-forming sequence from the insulin-linked polymorphic region.^36^ In contrast to the reduction in promoter activity for the c-Myc promoter in the presence of the peptides, there was no significant change in response in the insulin promoter (Fig.2A), demonstrating that Pep-VSE and Pep-PTN have some specificity for the iM from c-Myc.

**Figure 2.**
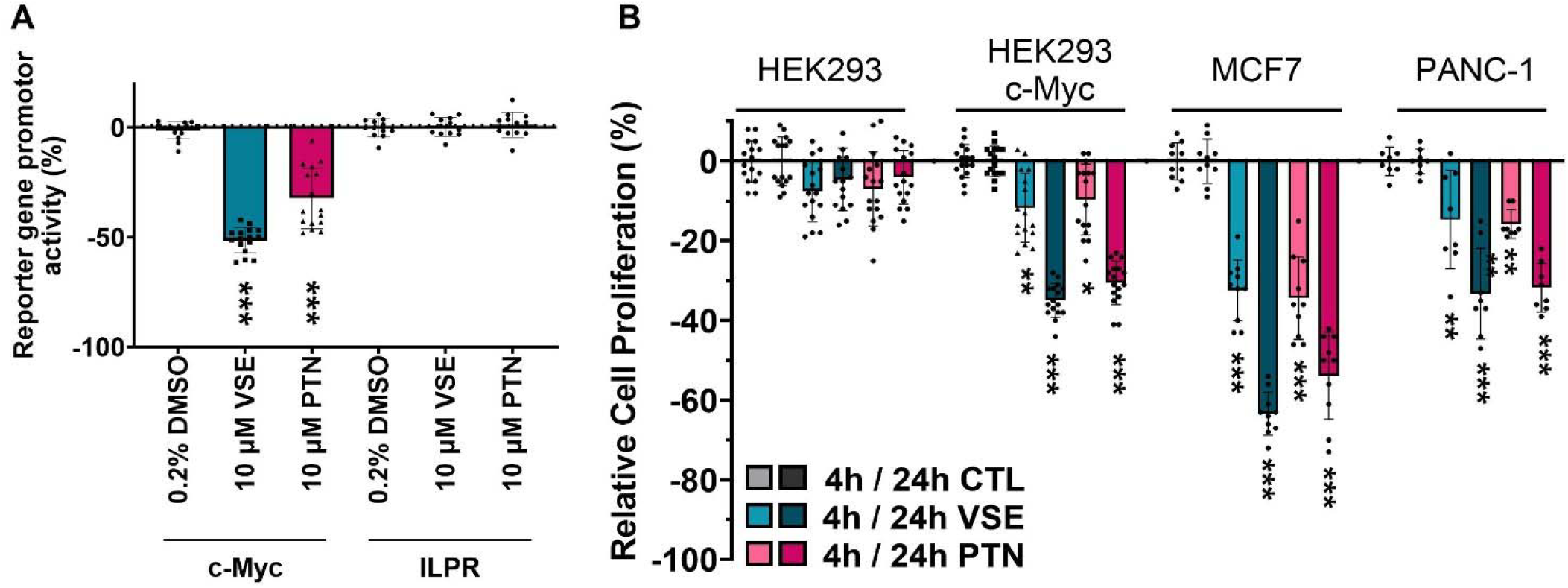
A) Difference in reporter gene activation in HEK293 cells transfected with c-Myc or ILPR after 4h of treatment in presence of Nanocin Pro^®^ (+) with peptide VSE or PTN at 10 µM. The scattered plots represent 8 biological repeats for HEK293(+c-Myc-reporter) and 6 biological repeats for HEK293(+ILPR-reporter) each with 2 technical repeats. Data shown are mean ± SD (n=5/6), *** p<0.001 determined by a one-way ANOVA with Holm - Šídák post-hoc analysis. B) Relative change in cell proliferation of HEK293, HEK293(+c-Myc-reporter), MCF7, and PANC-1 cells after 4h and 24h of treatment in presence of Nanocin Pro^®^ (+) with peptide VSE or PTN at 10 µM. The scattered plots represent 6 biological repeats for MCF-7 and 5 biological repeats for HEK293-ILPR each with 2 technical repeats. Data shown are mean ± SD (n=5/6), *** p<0.001 determined by a one-way ANOVA with Holm - Šídák post-hoc analysis.

Cell proliferation assays were conducted concurrently with the luciferase assays (Fig. 2B). The peptides had no significant effect on proliferation in HEK293 wildtype cells (Fig.2B and Fig.S15A). To examined whether there were any potential effects on cells which could have arisen from the reduction in c-Myc expression, we determined the effects on cell proliferation in MCF-7 and PANC-1 cells, examples of cell-lines in which proliferation is driven by c-Myc.^52, 53^ Both MCF-7 and PANC-1 cells showed significant time-dependent reduction in cell proliferation after addition of the peptides, in-line with an effect of suppressing c-Myc transcription firing on a background of high Myc-driven cell proliferation. Interestingly, we observed that HEK293 cells transfected with the c-Myc reporter gene, also demonstrated reduced proliferation in the presence of the peptides (Fig.2B), whereas the insulin reporter gene did not affect proliferation significantly (Fig.S15B). This is indicative of the diversion of the cellular machinery to the c-Myc reporter gene in HEK293 cells when exposed to the peptides.

Given the selectivity of the Pep-PTN and Pep-VSE peptides for the c-Myc iM structure, both in the binding experiments and also the *in vitro* gene assays, we considered potential binding modes. We recorded NMR spectra of the c-MycC52 oligonucleotide in the presence and absence of Pep-VSE to give insights into potential binding sites (Figs.S17-20). In the presence of one equivalent of Pep-VSE we observe no pertubations to imino resonances indicating that the DNA remains structured in the presence of peptide; we also observe that the peptide is soluble and not aggregated and the pH does not shift upon titration. The peptide spectra show some chemical shift changes in aromatic resonances (7-8 ppm, Fig.S19-20), which based on chemical shift and multiplicity are likely to arise from the tryptophan residues. The changes in signal are small which indicates either that binding occurs in the loops far from base pairs, hence we see no perturbation in these signals upon binding, or the peptide remains in a disordered conformation upon binding, so that chemical shifts are not perturbed far from their ensemble-averaged values upon binding. Given the specificity, binding and stabilization experiments indicating strong interactions between Pep-VSE and c-Myc, this is then consistent with a model where the peptide binds the loops, and not the core C-stack or grooves. The absence of structural information on the long c-Myc iM-forming sequence led us to create models based on the existing intramolecular iM structure from the insulin-linked polymorphic region^36^ and the previous folding conformations proposed by Sutherland *et. al*.^30^ (see supporting information). Molecular dynamics simulations were employed (Fig. S22-23) and subsequently docking was performed to explore interactions between the peptides and two distinct conformations of the c-Myc iM DNA. The highest-ranked model (characterised by the most negative docking score and highest confidence score) from each docking experiment was selected as the optimal peptide-DNA complex structure (Fig.3).

**Figure 3:**
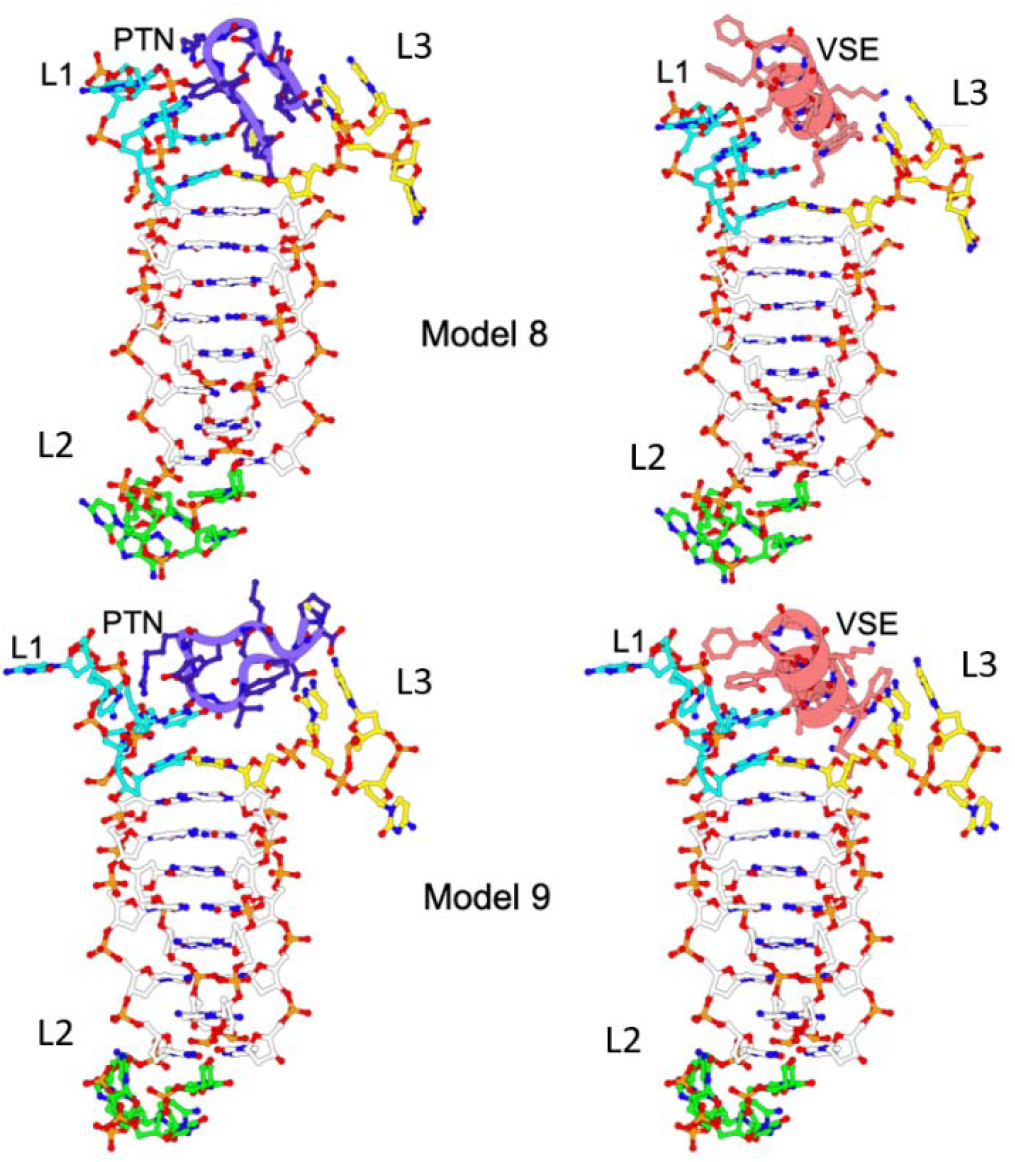
Models of the c-Myc iM and docked peptides Pep-PTN (purple) and Pep-VSE (salmon) between loops (L) (bases 10-14; cyan sticks) and 3 (bases 29-32; yellow sticks) in Model 8 (top row) and Model 9 (bottom row).

In this study we identified several peptides which target the c-MycC52 iM sequence. Two of which we have studied in more detail to reveal their selective binding for this iM-forming sequence and their selective modulation of c-Myc transcription in intact cells. Hurley and co-workers illuminated the mutually exclusive binding of nucleolin and hnRNP-K to the c-Myc G quadruplex and i-motif structures, and highlighted the challenge in the design of chemical probes and drug molecules that can precisely control c-Myc transcriptional firing.^30^ Aberrant activation of c-Myc transcription has the potential to further promote cancer cell growth and these peptides can serve as a counter-weight to this, either as probes on their own or in combination with other ligands. There is much potential in using peptides to target iMs. To date, the only other peptides that have been studied to target-iM structures were targeting Hif-1-α but were not assessed for their activity.^54^ The main benefit of these peptide ligands is their ease of use. Through custom commercially-available peptide synthesis there is great potential for their use as biological probes. These could be useful tools for studying c-Myc gene expression or may act as lead compounds for development of peptidomimetic therapeutics targeting c-Myc with enhanced drug-like properties. There are now multip e examples of successful development of small-molecule peptidomimetics,^55^ leaving this chemical-space now open for more-specific small molecules to be derived from these core peptide structures. Finally, through selecting these peptides by phage display we also demonstrate the potential for using phage display for other iM-targets in the human genome and beyond.

## Data Availability

All data will be available at 10.5281/zenodo.14258360.

## Acknowledgements

We thank Dr Nikita Harvey for supporting the NMR experiments, which were acquired using the UCL School of Pharmacy NMR Core Facility (RRID:SCR_027123).

## Author Contributions

Z.A.E.W. and C.J.M. secured the research funding, conceived and designed the study; S.R. performed and analysed the Phage Display and FID experiments; D.G. designed, performed and analysed the stabilization and binding experiments and all cell-based studies; J.K. Supported D.G. with some of the preliminary reporter gene assays. E.A., S.C. and S.H. designed, performed and analysed the computational experiments. C.A.W. Performed and analysed the NMR experiments. S.R., D.G., C.J.M. and Z.A.E.W. wrote the first draft of the paper. S.R., D.G., E.A., S.H., C.W., C.J.M. and Z.A.E.W. contributed to the review and editing of the manuscript.

