## Supplementary data for "Affinity-selected peptide ligands specifically bind i-motif DNA and modulate c-Myc gene expression"

### 1) Oligonucleotide Sequences

**Table S1.** Target oligonucleotide sequences. Where the text refers to B-DNA or dsDNA this consists of the ds and ds comp sequences annealed together.

| *Oligonucleotide* | *Sequence (5' - 3')* |
|---|---|
| *c-MycC27* | CCT-TCC-CCA-CCC-TCC-CCA-CCC-TCC-CCA |
| *c-MycC52* | CTT-CTC-CCC-ACC-TTC-CCC-ACC-CTC-CCC-ACC-CTC-CCC-ATA-AGC-GCC-CCT-CCC-G |
| *c-MycG27* | TGG-GGA-GGG-TGG-GGA-GGG-TGG-GGA-AGG |
| *c-MycG52* | CGG-GAG-GGG-CGC-TTA-TGG-GGA-GGG-TGG-GGA-GGG-TGG-GGA-AGG-TGG-GGA-GAA-G |
| *ds* | GGC-ATA-GTG-CGT-GGG-CGT-TAG-C |
| *ds comp* | GCT-AAC-GCC-CAC-GCA-CTA-TGC-C |
| *DAPc* | CCC-CCG-CCC-CCG-CCC-CCG-CCC-CCG-CCC-CC |
| *DAPg* | GGG-GGC-GGG-GGC-GGG-GGC-GGG-GGC-GGG-GG |

**Table S2.** Competitor oligonucleotide sequences used in phage display selection rounds. Where dsDNA consists of the ds and ds comp sequences annealed and the Holliday Junction consists of the Hjb, Hjh, Hjr, and Hjx annealed together.

| *Oligonucleotide* | *Sequence (5' - 3')* |
|---|---|
| *ATXN2L* | CCC-CCC-CCC-CCC-CCC-CCC-CCC-CCC |
| *C-hairpin* | CTC-TCT-TCT-CTT-CAT-TTT-TCA-ACA-CAA-CAC-AC |
| *DAP* | CCC-CCG-CCC-CCG-CCC-CCG-CCC-CCG-CCC-CC |
| *ds* | GGC-ATA-GTG-CGT-GGG-CGT-TAG-C |
| *ds comp* | GCT-AAC-GCC-CAC-GCA-CTA-TGC-C |
| *Hif1$\alpha$* | CGC-GCT-CCC-GCC-CCC-TCT-CCC-CTC-CCC-GCG-C |
| *hTeloC* | TAA-CCC-TAA-CCC-TAA-CCC-TAA-CCC |
| *hTeloG* | GGG-TTA-GGG-TTA-GGG-TTA-GGG |
| *ILPR* | TGT-CCC-CAC-ACC-CCT-GTC-CCC-ACA-CCC-CTG-T |
| *NasC* | GGG-CGG-GCT-GGG-CAT-TGC-GGG |
| *NasT* | GGG-AGC-GGG-ACG-GGG-GCC-GGG |
| *Hjb* | CGG-TAG-CAG-TAC-CGT-TGG-TGG-C |
| *Hjh* | GCC-TAG-CAT-GAT-ACT-GCT-ACC-G |
| *Hjr* | GCC-ACC-ACC-GGC-GTC-AAC-TGC-C |
| *Hjx* | GGC-AGT-TGA-CGT-CAT-GCT-AGG-C |
| *RNA C1UUU* | CUU-UCU-UUC-UUU-CUU-UC |

### 2) Biophysical Characterisation of cMycC52 iM-Forming Sequence

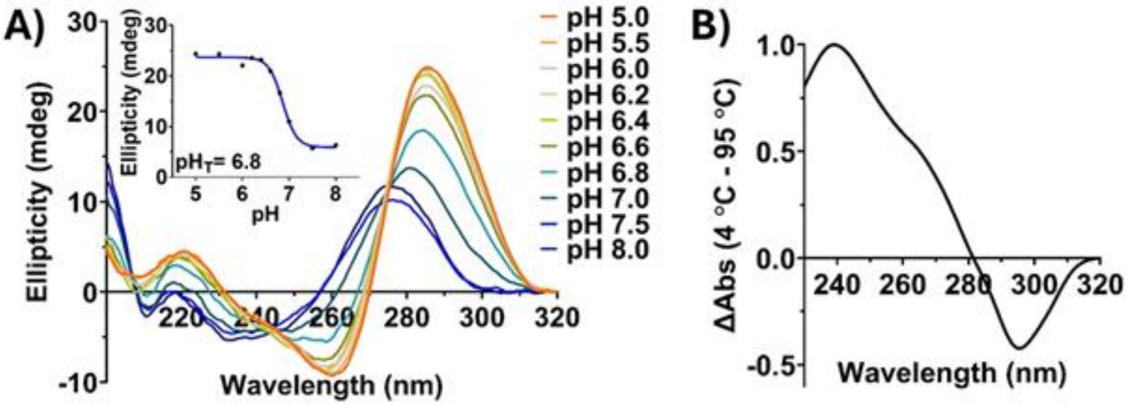

**Figure S1.** (A) Circular dichroism (CD) spectrum of 10 µM c-MycC52 in 10 mM sodium cacodylate buffer across a pH range of 5.0 to 8.0 to show pH-dependent i-motif structure formation. The inset shows the determination of the transitional pH, obtained by plotting the ellipticity at 288 nm against pH. The transitional pH corresponds to the condition at which 50% of the DNA population is folded into the i-motif. (B) Thermal difference spectra of c-MycC52 with 2.5 µM DNA in 10 mM NaCaco at pH 6.6. Source data for this figure are provided as a Source Data file.

### 3) Phage Display and Peptides

**Table S3.** Phage display methods for panning against c-MycC52 using the NEB Ph.D.TM12 linear randomized 12-mer library.

| *Round* | *Phage* | *Blocking buffer* | *Incubation Buffer* | *Wash buffer* | *Elution Buffer* | *Concentration of 5'-biotinylated c-MycC52 (pmol)* | *Non-biotinylated competitors added to phage mixture* | *Output titer (pfu)* |
|---|---|---|---|---|---|---|---|---|
| *1* | $1x10^{11}$ Phage Library, Ph.D.-12, in 500 μL PBS pH 6.0 | PBS pH 6.0, 5% BSA | PBS pH 6.0 | PBS pH 6.0, 0.1% Tween 20 | PBS, pH 7.4 | 10 | None | $1.72 \times 10^6$ |
| *2* | $1x10^{11}$ Amplified Output Phage from Round 1, Elution 3 in 500 μL PBS pH 6.0 | PBS pH 6.0, 5% BSA | PBS pH 6.0 | PBS pH 6.0, 0.5% Tween20 | PBS, pH 7.4 | 1 | 100 pmol of each competitor: NasT, NasC, hTeloG, C-hairpin, Holiday Junction, dsDNA, Calf Thymus, RNA C1UUU | $5.11 \times 10^6$ |
| *3* | $1x10^{11}$ Amplified Phage from Panning 2, Elution 3 in 500 μL PBS pH 6.0 | PBS pH 6.0, 5% BSA | PBS pH 6.0 | PBS pH 6.0, 0.5% Tween20 | PBS, pH 7.4 | 1 | 100 pmol of each competitor: hTeloC, ILPR, ATXN2L, DAP, Hif1α | $2.52 \times 10^7$ |

**Table S4.** Sequences, structures, and purity (determined by RP-HPLC) of the peptides used in this study

| *Peptide* | *Sequence* | *Structure* | *Purity %* |
|---|---|---|---|
| *SLC* | SLCDIIRIEKVR | 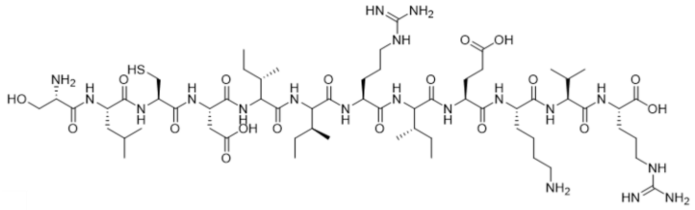 | 95.45 |
| *PTN* | PTNVSGRYLFC | 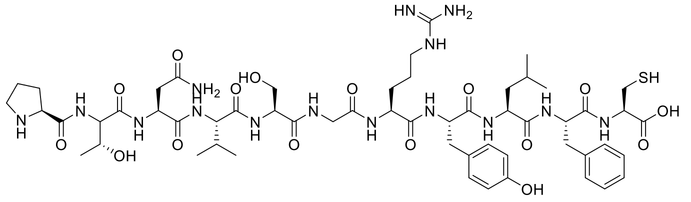 | 95.09 |
| *VSE* | VSEAWKEVKGFF | 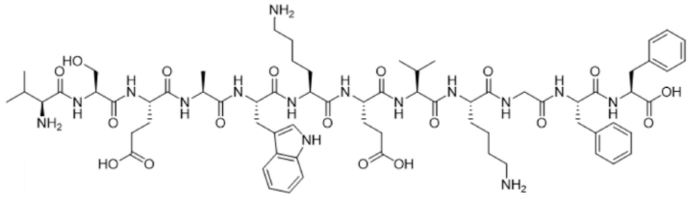 | 98.49 |
| *EIE* | EIEYTDHMKELG | 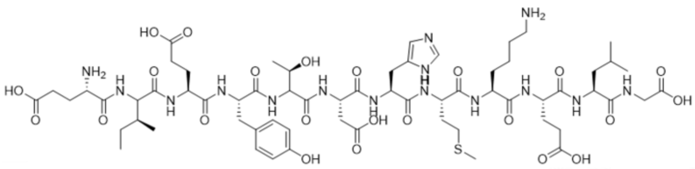 | 96.10 |
| *RVS* | RVSTDHMKGRGG | 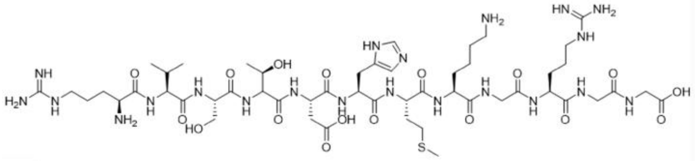 | 95.70 |

### 4) Circular Dichroism Melting Experiments

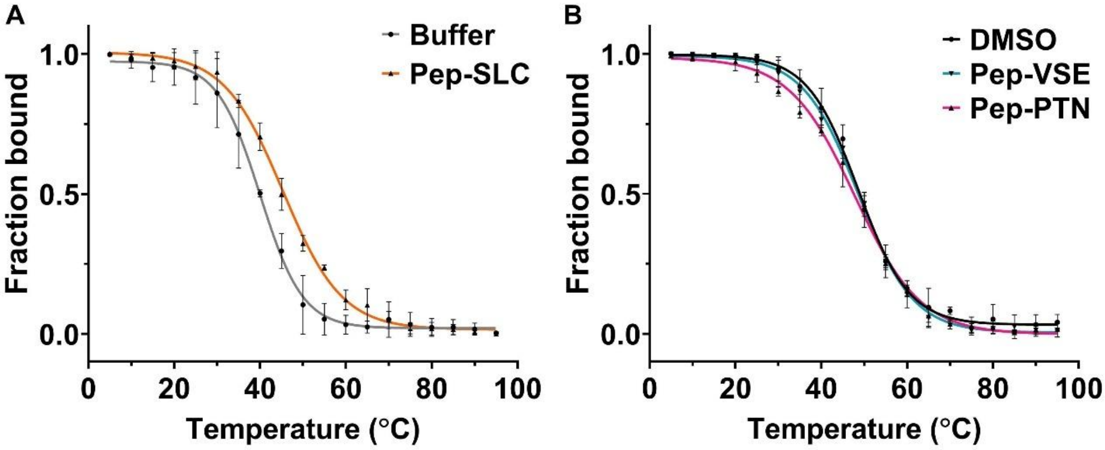

**Figure S2.** Normalised fraction folded taken from ellipticity at 253 nm 10 µM DS in presence of 10 mM NaCaco at pH 6.6 buffer and 10 molar eqv. Pep-SLC (A) and in presence of 10 molar eqv. DMSO, Pep-VSE, or Pep-PTN. All Data are presented as Mean ± SD (n=2) and a sigmoidal fitting.

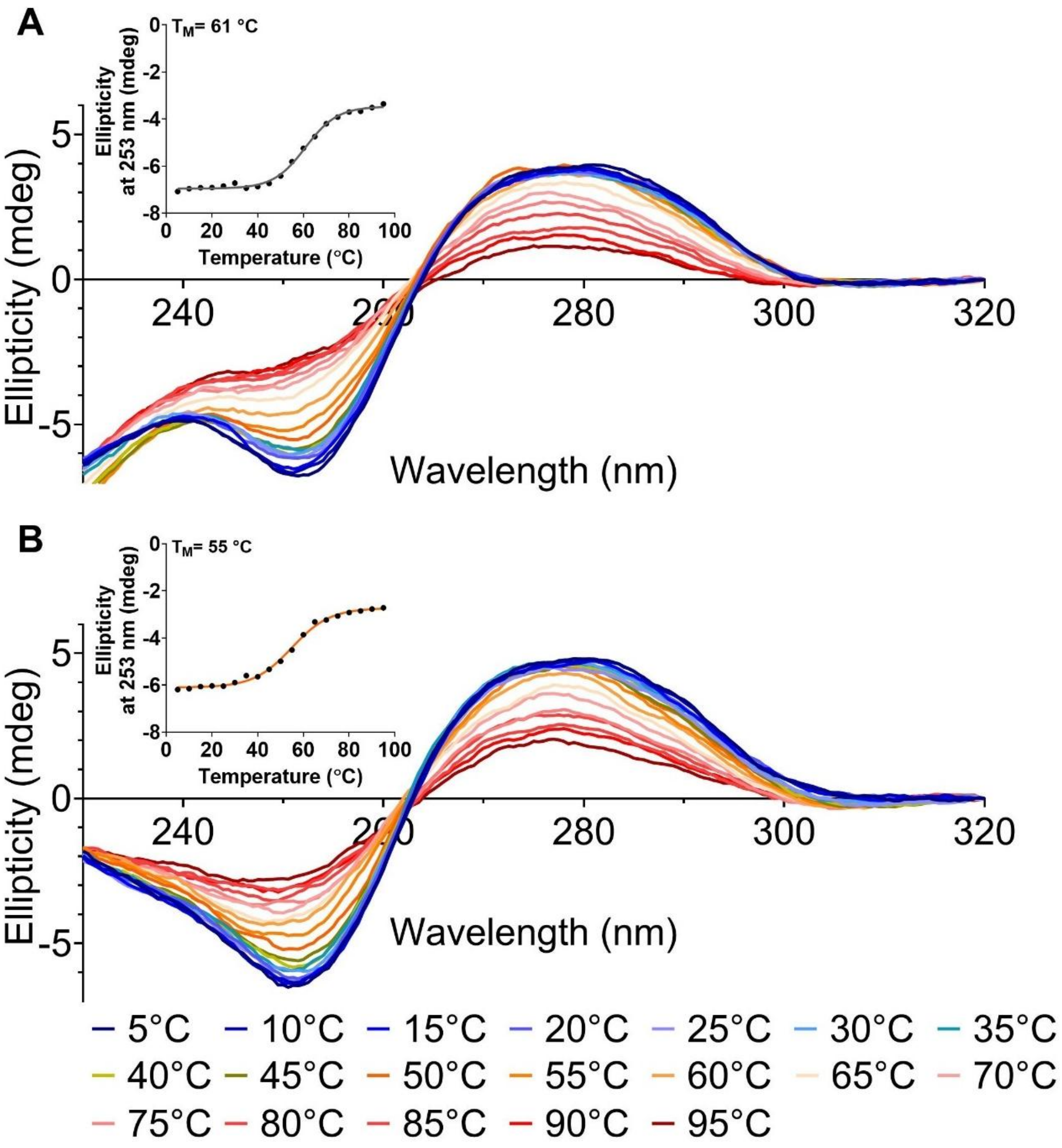

**Figure S3.** Example CD melting experiments with 10 μM DS in 10 mM NaCaco pH 6.6 buffer (A) and presence of 10 molar eqv. SLC (B) with inserts showing corresponding sigmoidal curve fitting model for ellipticity vs temperature.

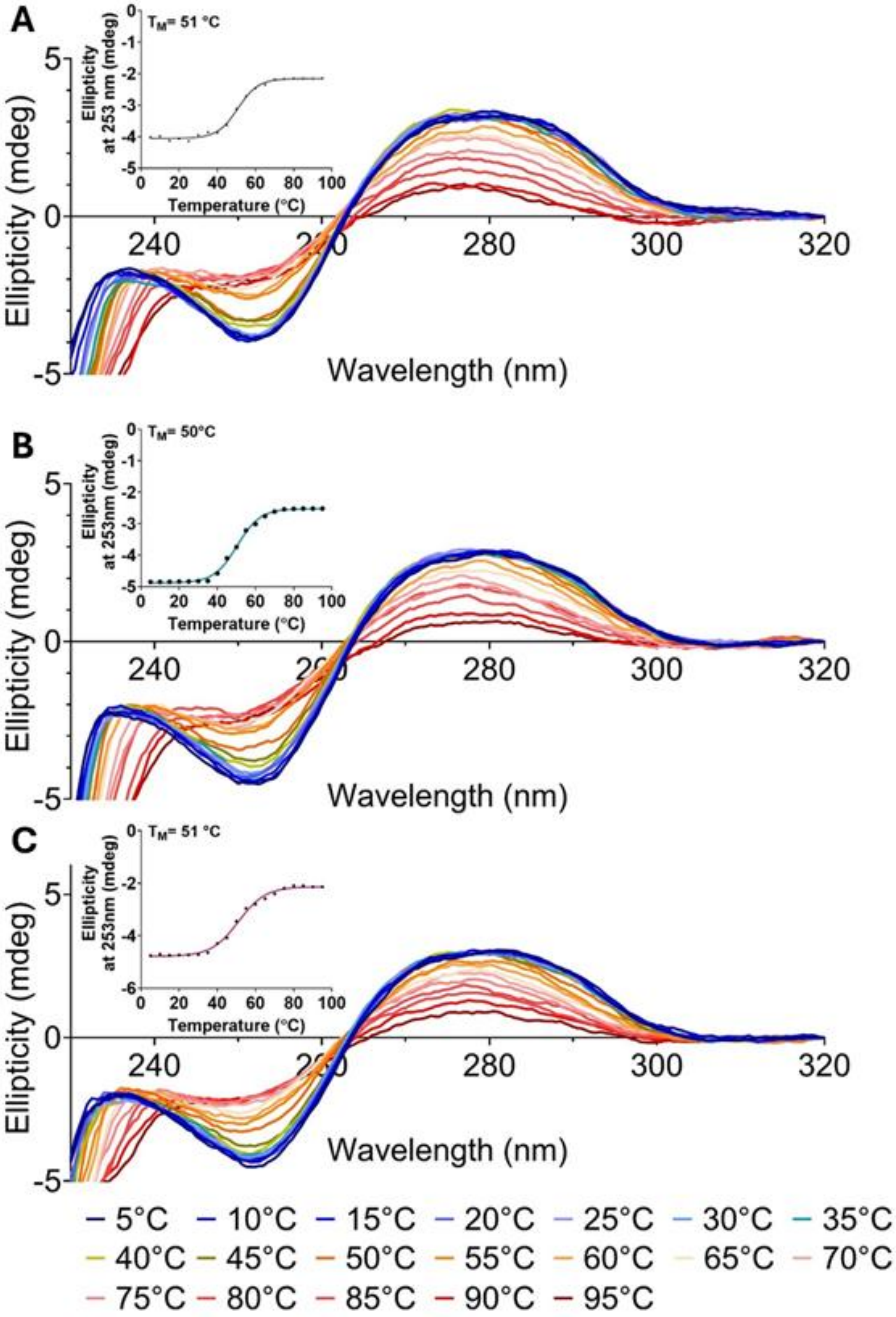

**Figure S4.** Example CD melting experiments with 10 μM DS in 10 mM NaCaco pH 6.6 buffer and presence of 10 molar eqv. DMSO (A), Pep-VSE (B) or Pep-PTN (C). Inserts showing corresponding sigmoidal curve fitting model for ellipticity vs temperature.

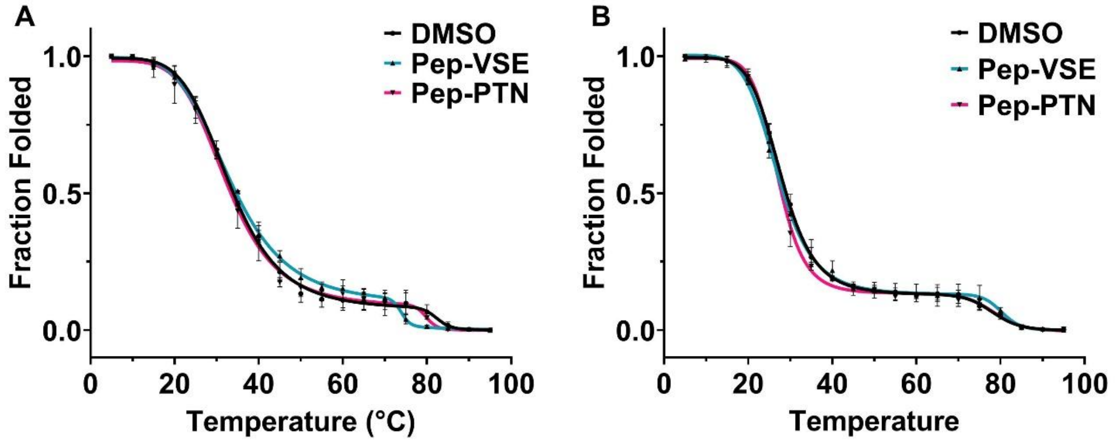

**Figure S5.** Normalised fraction folded taken from ellipticity at 288 nm 10 µM cMycC52 (A) and cMycC27 (B) in 10 mM NaCaco at pH 6.6 buffer in presence of in the presence of 10 molar eqv. DMSO, Pep-PTN or Pep-VSE. All Data are presented as Mean ± SD (n=3).

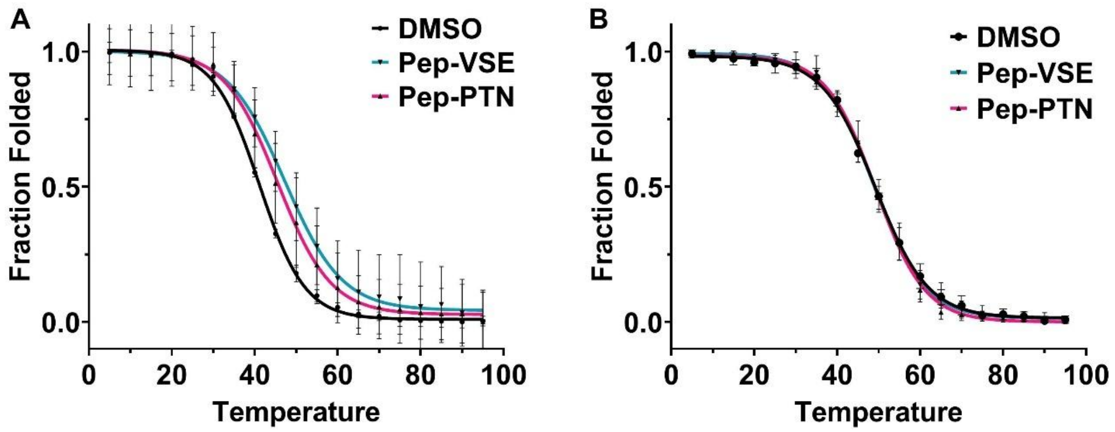

**Figure S6.** Normalised fraction folded taken from ellipticity at 264 nm 10 µM cMycG52 (A) and cMycG27 (B) in 10 mM NaCaco at pH 6.6 buffer in presence of in the presence of 10 molar eqv. DMSO, Pep-PTN or Pep-VSE. All Data are presented as Mean ± SD (n=3).

**Table S5.** CD melting analysis of 10 µM c-MycC27 and c-MycC52 and their complimentary c-MycG sequences in 10 mM NaCaco at pH 6.6 buffer in the presence of 10 molar eqv. DMSO, Pep-PTN or Pep-VSE. All Data are presented as Mean ± SEM (n=4) and statistical significance via One-way ANOVA with Bonferroni post-hoc analysis is expressed in bold as ns>0.1, p<0.05*, p<0.01**, p<0.001***.

| | $T_M$ (°C) DMSO | $T_M$ (°C) PTN | $T_M$ (°C) VSE |
|---|---|---|---|
| *c-MycC27* | 27 ± 0.8<br>79 ± 1.0 | 27 ± 0.7<br>79 ± 0.5 | 26 ± 1.0<br>80 ± 0.6 |
| *c-MycC52* | 33 ± 0.8<br>83 ± 0.9 | 32 ± 2.0<br>79 ± 0.8 | 33 ± 0.9<br>74 ± 0.7 |
| *c-MycG27* | 49 ± 0.5 | 50 ± 1.0 | 48 ± 1.0 |
| *c-MycG52* | 41 ± 0.1 | 45 ± 1.0 | 48 ± 1.0 |
| *DS* | 51 ± 0.9 | 52 ± 1.0 | 51 ± 0.8 |

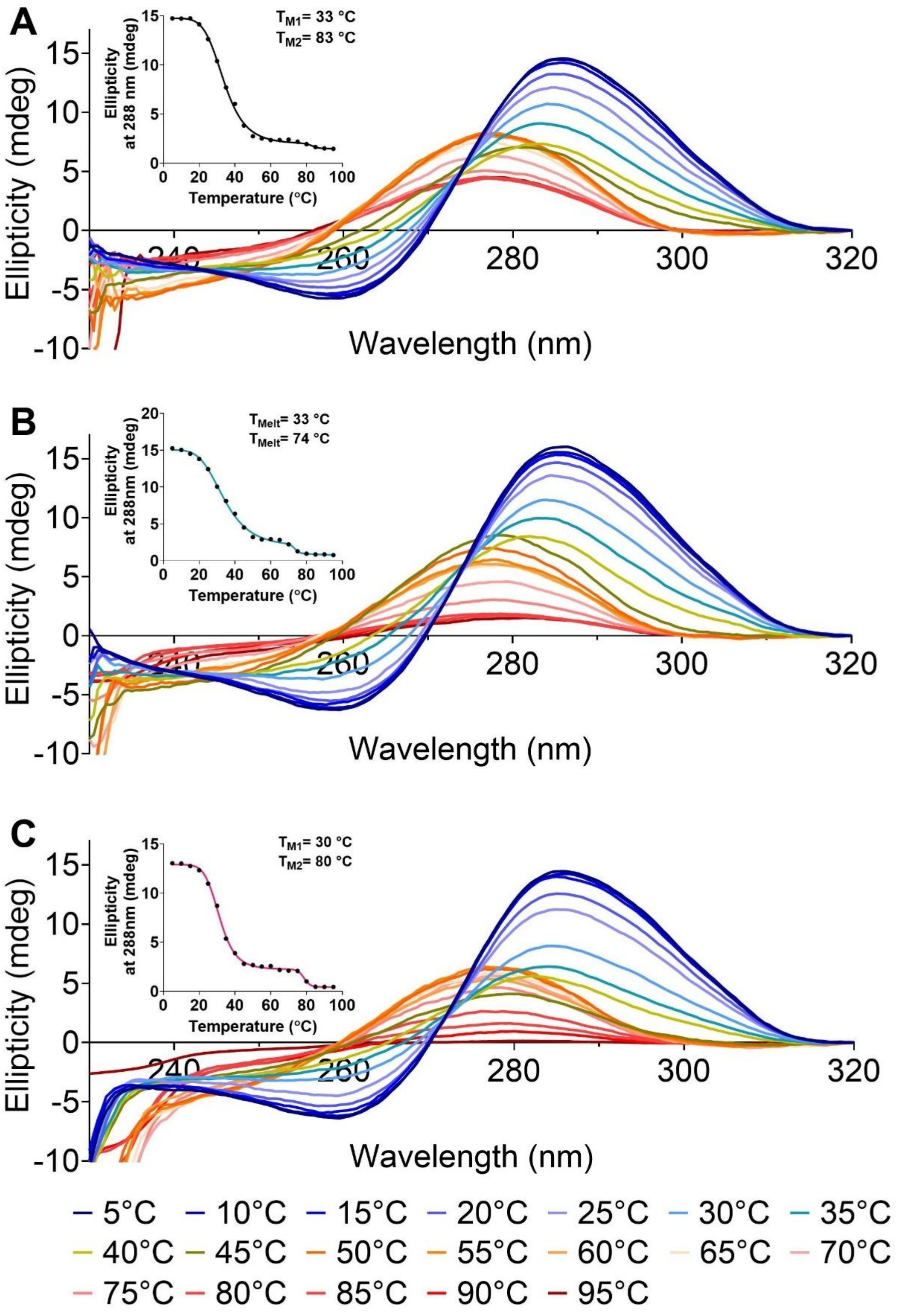

**Figure S7.** Example CD melting experiments with 10 µM c-MycC52 in 10 mM NaCaco pH 6.6 buffer and presence of 10 molar eqv. DMSO (A), Pep-VSE (B) or Pep-PTN (C). Inserts showing corresponding biphasic sigmoidal curve fitting model for ellipticity vs temperature.

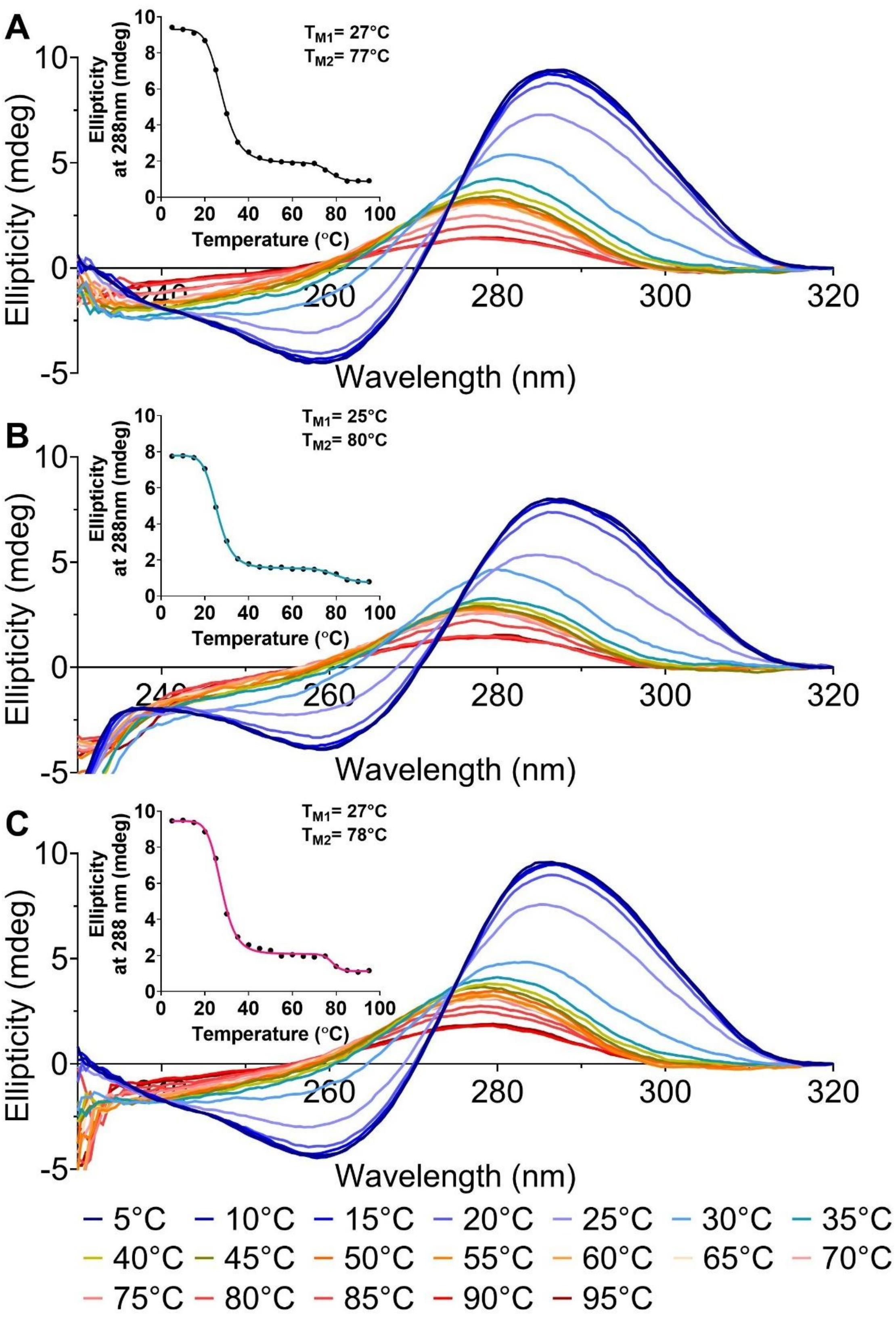

**Figure S8.** Example CD melting experiments with 10 µM c-MycC27 in 10 mM NaCaco pH 6.6 buffer and presence of 10 molar eqv. DMSO (A), Pep-VSE (B) or Pep-PTN (C). Inserts showing corresponding biphasic sigmoidal curve fitting model for ellipticity vs temperature.

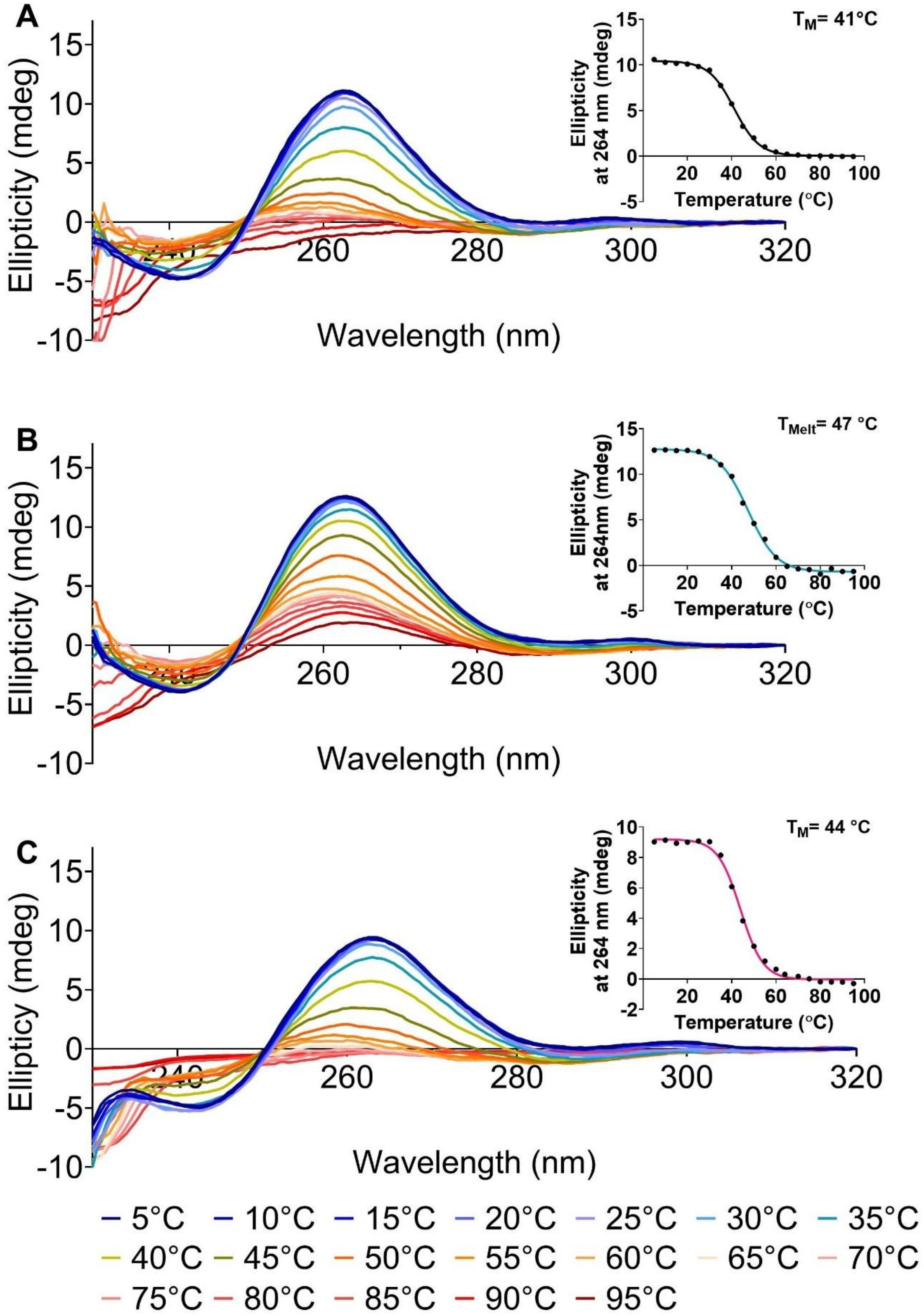

**Figure S9.** Example CD melting experiments with 10 µM c-MycG52 in 10 mM NaCaco pH 6.6 buffer and presence of 10 molar eqv. DMSO (A), Pep-VSE (B) or Pep-PTN (C). Inserts showing corresponding sigmoidal curve fitting model for ellipticity vs temperature.

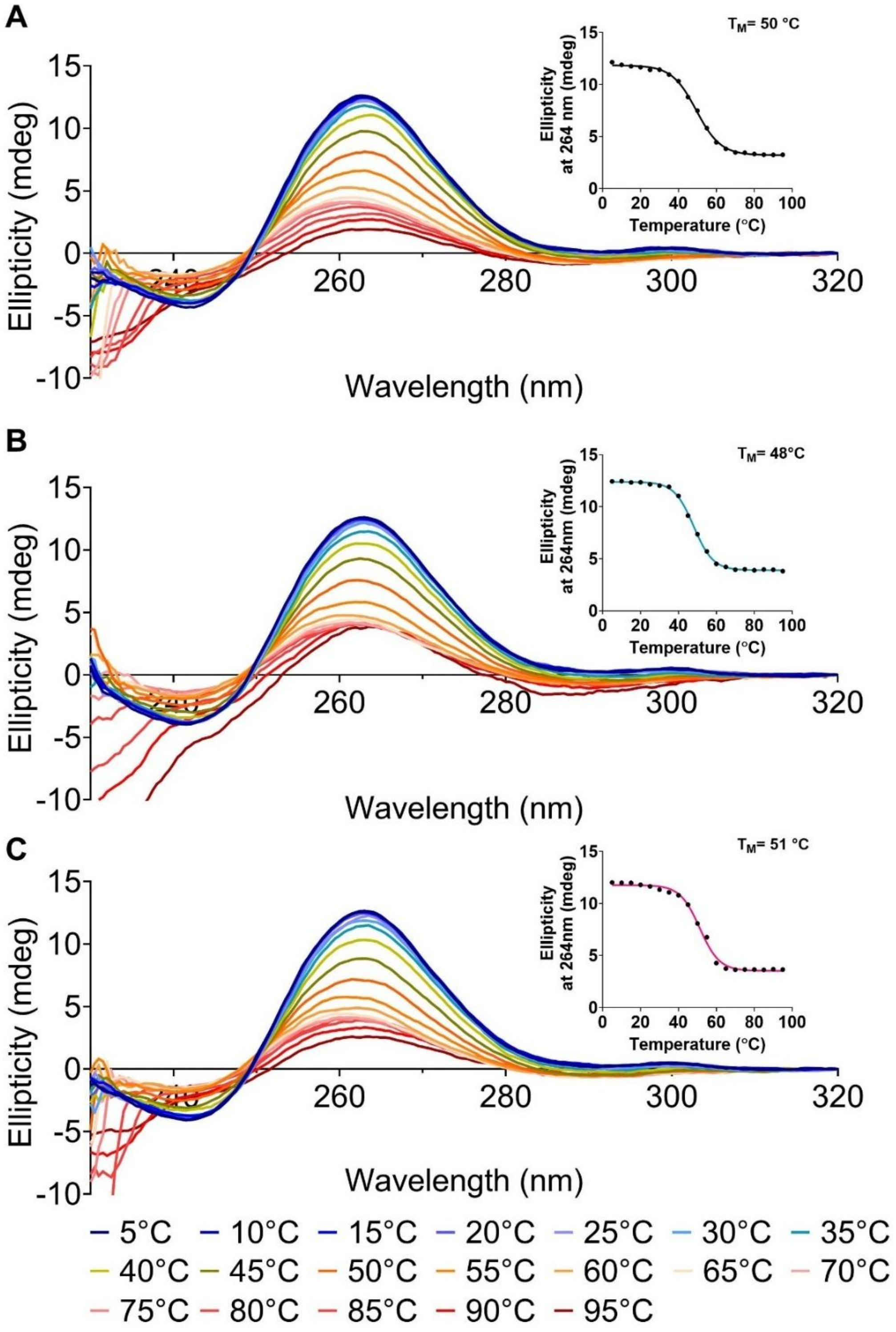

**Figure S10.** Example CD melting experiments with 10 µM c-MycG27 in 10 mM NaCaco pH 6.6 buffer and presence of 10 molar eqv. DMSO (A), Pep-VSE (B) or Pep-PTN (C). Inserts showing corresponding sigmoidal curve fitting model for ellipticity vs temperature.

### 5) UV-Titration Experiments

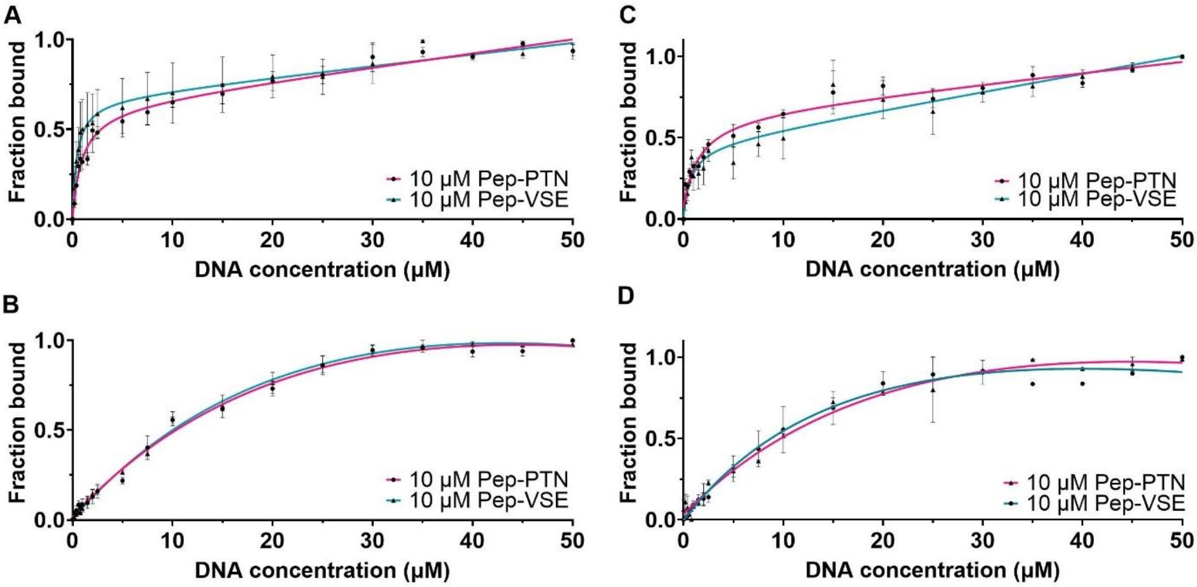

**Figure S11.** Normalised UV absorbance representing fraction bound of cMyc-C52 (A,C) and cMyc-G52 (B,D) obtained at absorbance maximum at 350 nm of 10 μM Pep-PTN or 10 μM Pep-VSE in 10 mM sodium cacodylate at pH 6.6 (A,B) and in presence of 100 mM potassium chloride (C, D). DNA was annealed in the same buffer and added as 0-50 μM DNA concentration. Data shown as Mean ± SD, n=3 and fitted with one site – total binding model in GraphPad.

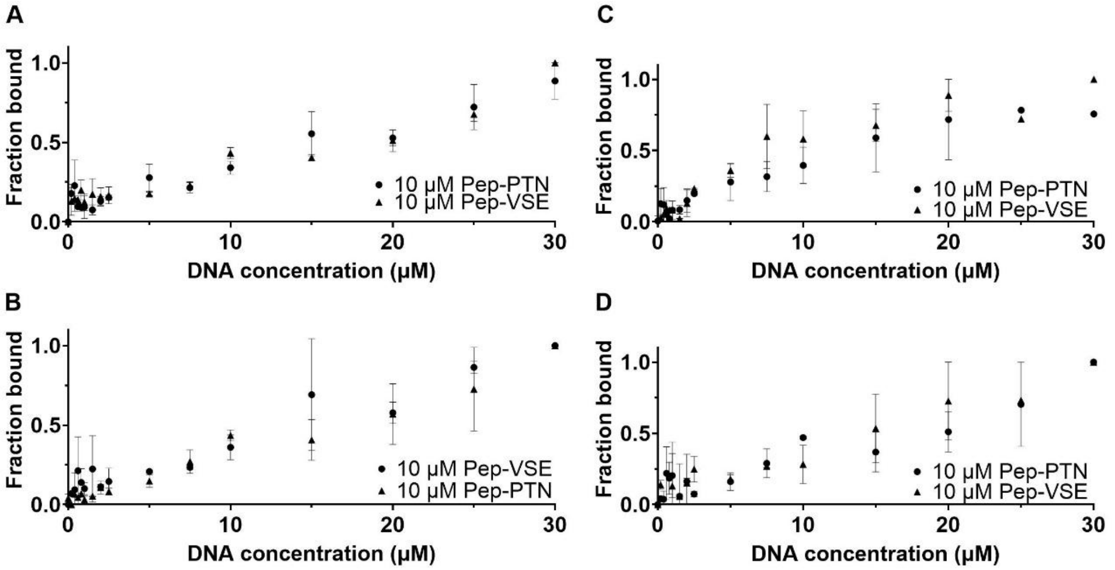

**Figure S12.** Normalised UV absorbance representing fraction bound of DAPc (A,C) and DAPg (B,D) obtained at absorbance maximum at 350 nm of 10 μM Pep-PTN or 10 μM Pep-VSE in 10 mM sodium cacodylate at pH 6.6 (A,B) and in presence of 100 mM potassium chloride (C, D). DNA was annealed in the same buffer and added as 0-30 μM DNA concentration. Data shown as Mean ± SD, n=2.

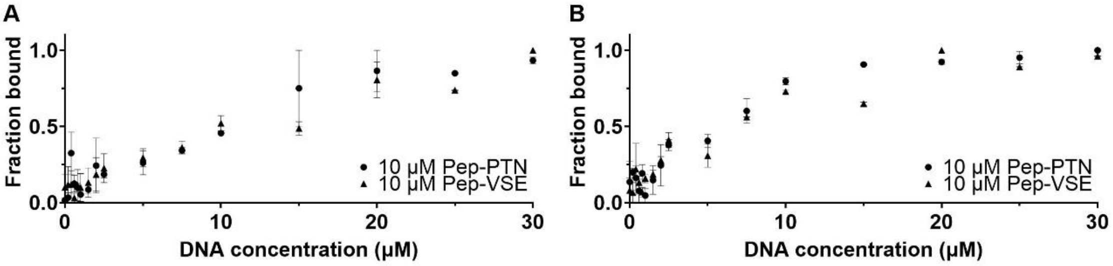

**Figure S13.** Normalised UV absorbance representing fraction bound of dsDNA obtained at absorbance maximum at 350 nm of 10 μM Pep-PTN or 10 μM Pep-VSE in 10 mM sodium cacodylate at pH 6.6 (A) and in presence of 100 mM potassium chloride (B). DNA was annealed in the same buffer and added as 0-50 μM DNA concentration. Data shown as Mean ± SD, n=2.

**Table S6.** 250 μM DNA annealed in 10 mM sodium cacodylate 100 mM potassium chloride at pH 6.6. Values represent the average of three experiments for cMyc sequences and two experiments for DAP and dsDNA sequences and the errors are standard deviations. NS =. Binding classified as non-specific (NS) showed no consistent or concentration-dependent pattern, under the tested conditions.

| Ligand | $K_d$ (μM) | | | | |
|---|---|---|---|---|---|
| | c-MycC52 | c-MycG52 | DAPc | DAPg | dsDNA |
| PTN | 1.54 ± 0.03 | 45 ± 4 | NS | NS | NS |
| VSE | 0.69 ± 0.06 | 35 ± 3 | NS | NS | NS |

### 6) Cell Experiments

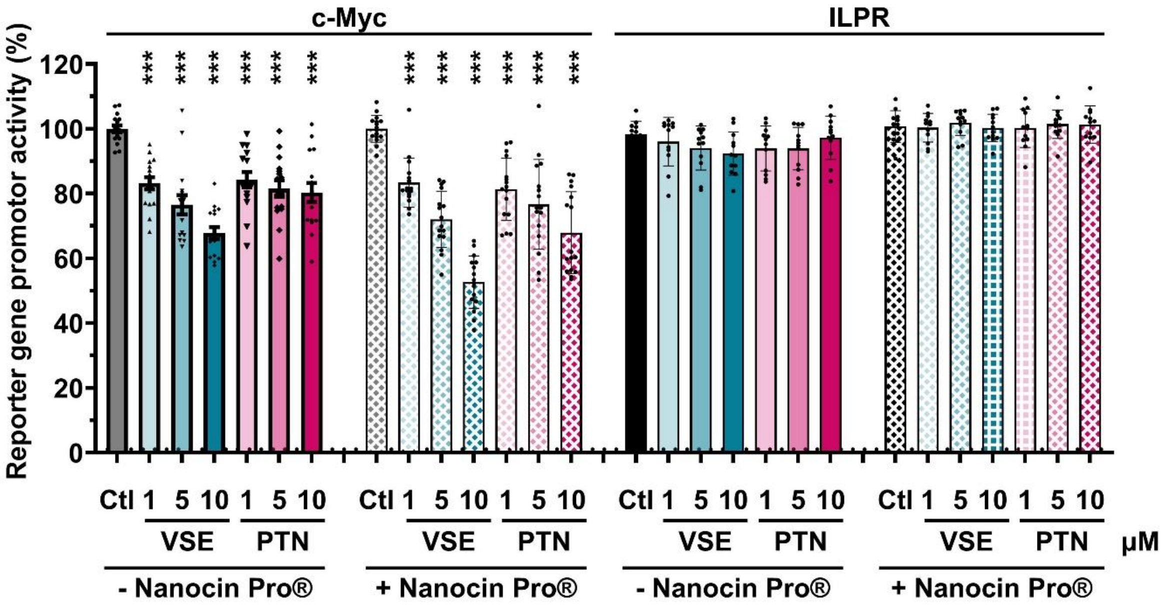

**Figure S14**. Reporter gene assay of HEK293 cells transfected with firefly luciferase regulated by the human promotor regions of c-Myc or ILPR after 4h of treatment in absence of Nanocin Pro® (-) and presence of Nanocin Pro® (+) with peptide VSE or PTN at final concentrations of 1 µM, 5 µM and 10 µM. The scattered plots represent 8 biological repeats for HEK293 – c-Myc and 6 biological repeats for HEK293 -ILPR each with 2 technical repeats. Data shown are mean ± SD (n=8/6), *** $p<0.001$ determined by a one-way ANOVA with Holm - Šídák post-hoc analysis.

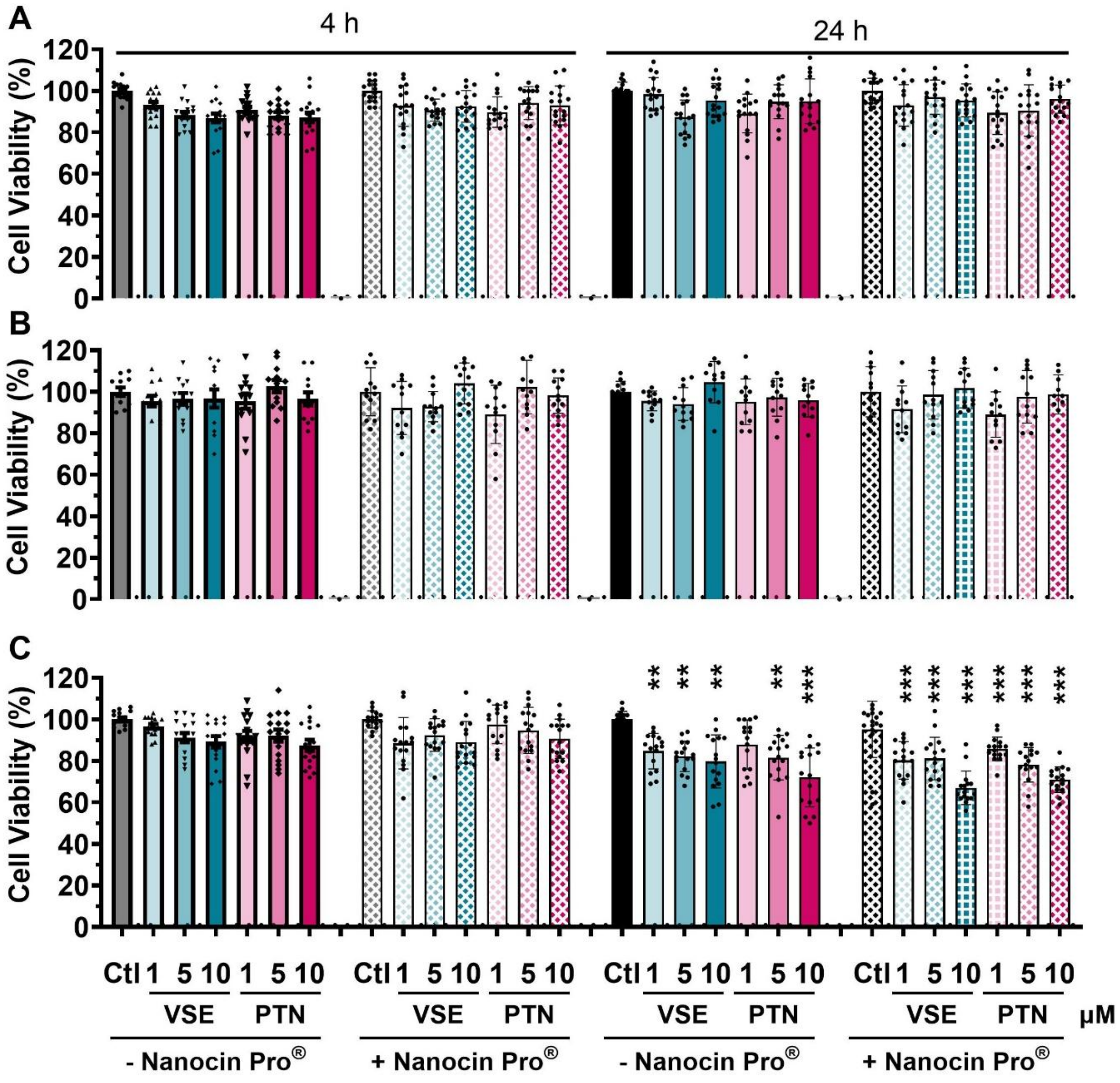

**Figure S15**. Cell viability of HEK293 wildtype (A), HEK293 – ILPR (B), HEK293 – cMyc (C) after 4h and 24h of treatment in absence of Nanocin Pro® (-) and presence of Nanocin Pro® (+) with peptide VSE or PTN at final concentrations of 1 µM, 5 µM and 10 µM. The scattered plots represent 8 biological repeats for HEK293 WT and HEK293 – cMyc and 6 biological repeats for HEK293 -ILPR each with 2 technical repeats. Data shown are mean ± SD (n=8/6), ** $p<0.01$, *** $p<0.001$ determined by a one-way ANOVA with Holm - Šídák post-hoc analysis.

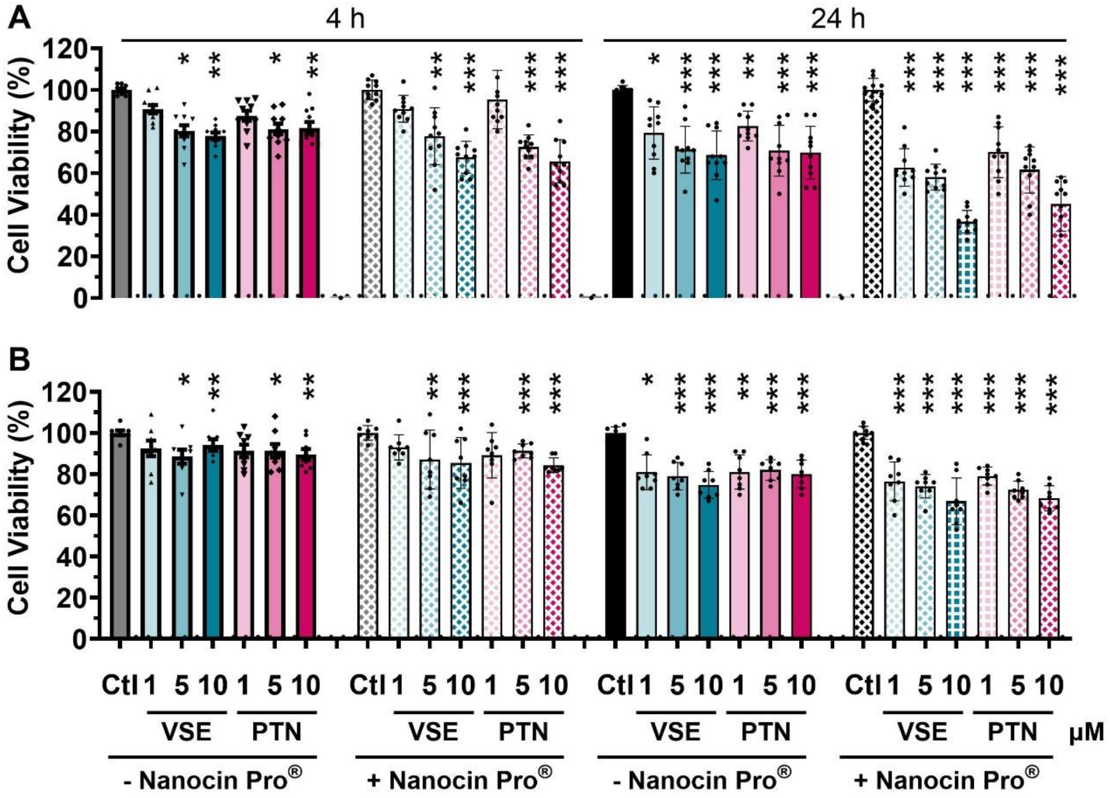

**Figure S16.** Cell viability of MCF7 (A) and PANC-1 cells (B) after 4h and 24h of treatment in absence of Nanocin Pro® (-) and presence of Nanocin Pro® (+) with peptide VSE or PTN at final concentrations of 1 μM, 5 μM and 10 μM. The scattered plots represent 6 biological repeats for MCF-7 and 4 biological repeats for Panc-1 cells each with 2 technical repeats. Data shown are mean ± SD (n=5/4), * $p<0.05$ ** $p<0.01$ *** $p<0.001$ determined by a one-way ANOVA with Holm - Šídák post-hoc analysis.

### 7) NMR Experiments

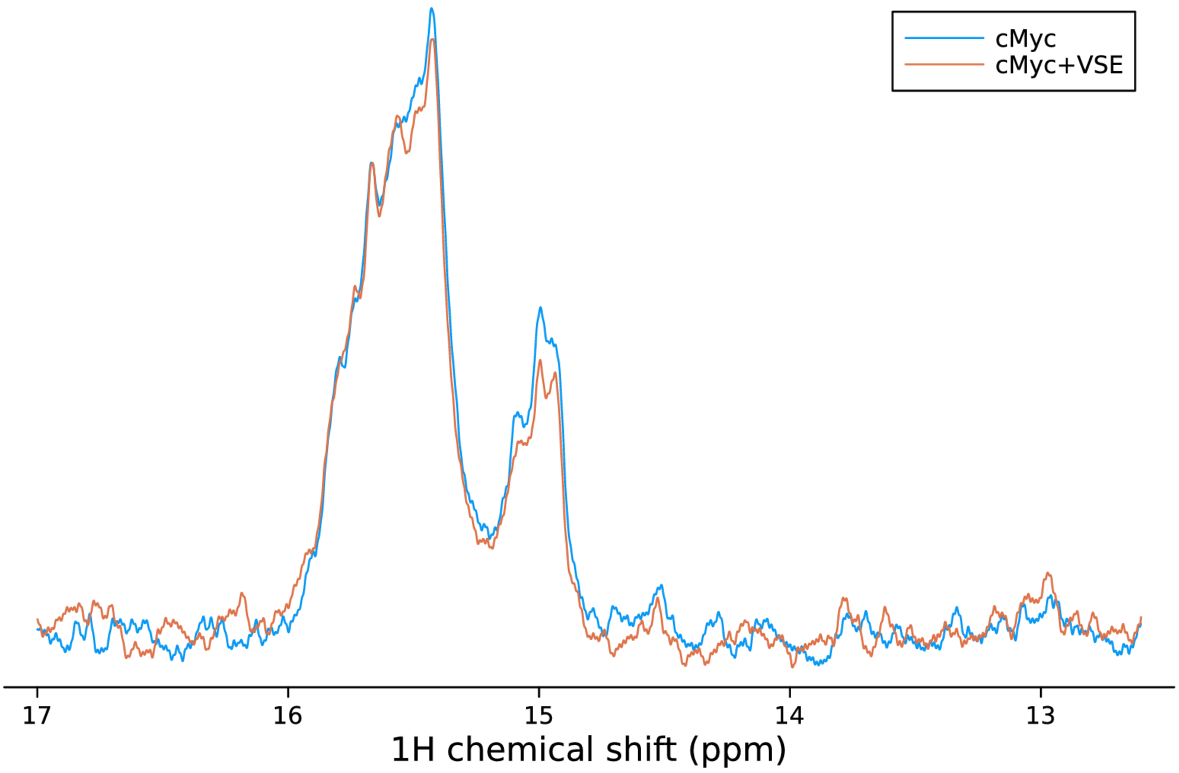

**Figure S17:** $^{1}H$ SOFAST-1D NMR spectra (600 MHz, 283 K) showing the imino region of 50 μM c-MycC52 and c-MycC52+Pep-VSE (1 eq) in 10 mM NaCaco, pH 6.6, 10% $D_2O$.

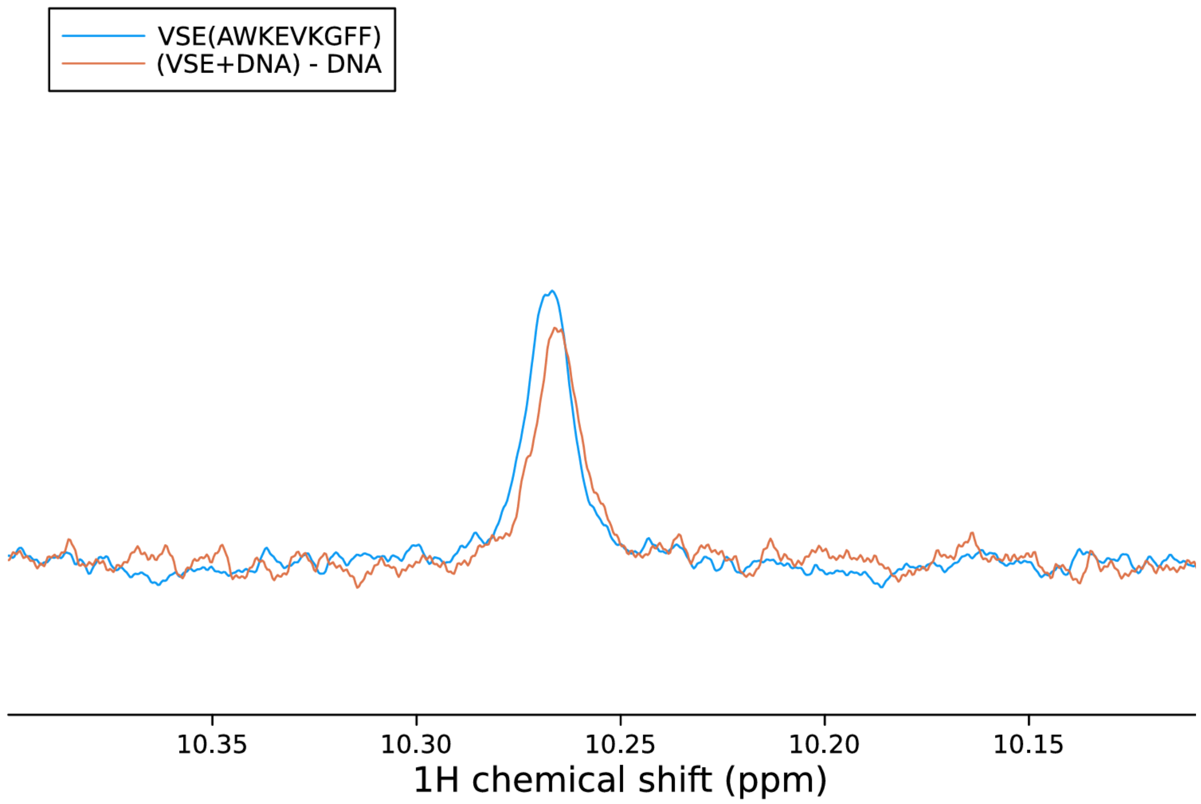

**Figure S18:** $^{1}$H 1D NMR spectra (600 MHz, 283 K) of Pep-VSE (50 μM, 10 mM NaCaco, pH 6.6, 10% $D_2O$) showing the tryptophan indole resonance (blue line); a comparison in the presence of 1 equivalent c-MycC52 is shown in red (an equivalent spectrum of DNA alone has been subtracted for clarity).

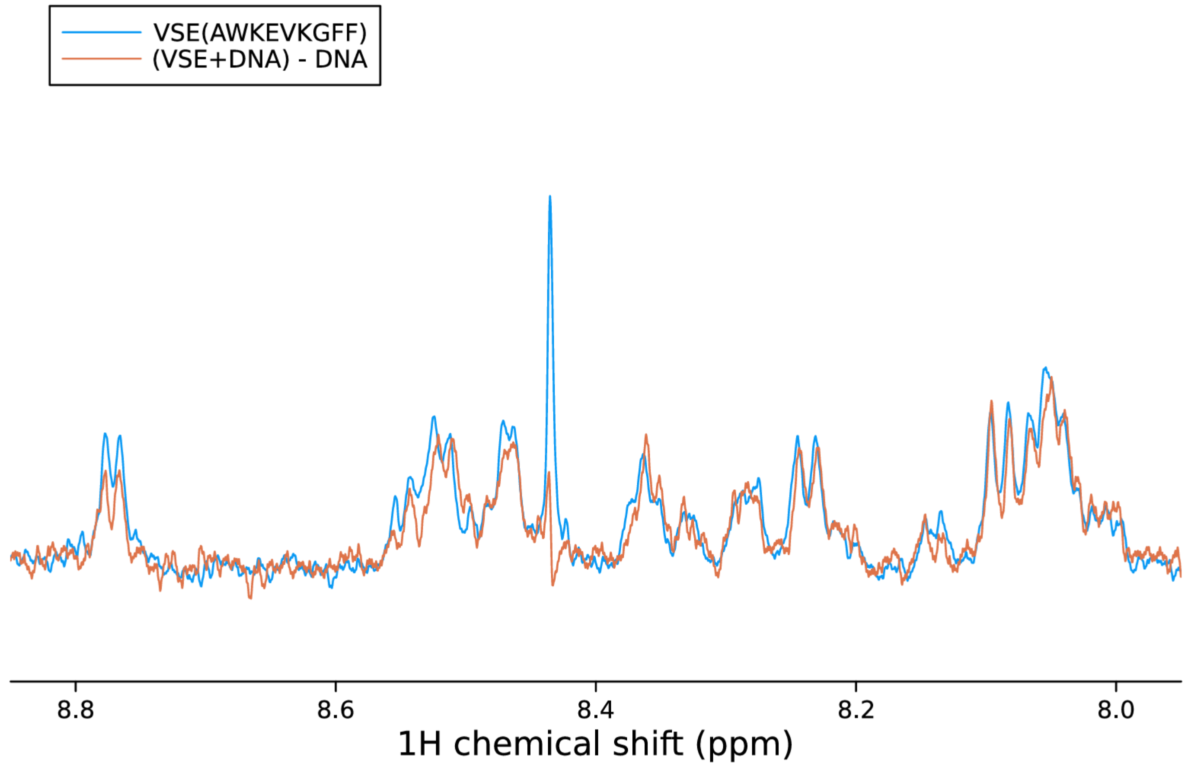

**Figure S19:** $^{1}$H 1D NMR spectra (600 MHz, 283 K) of Pep-VSE (50 μM, 10 mM NaCaco, pH 6.6, 10% $D_2O$) showing the amide resonances (blue line); a comparison in the presence of 1 equivalent c-MycC52 is shown in red (an equivalent spectrum of DNA alone has been subtracted for clarity).

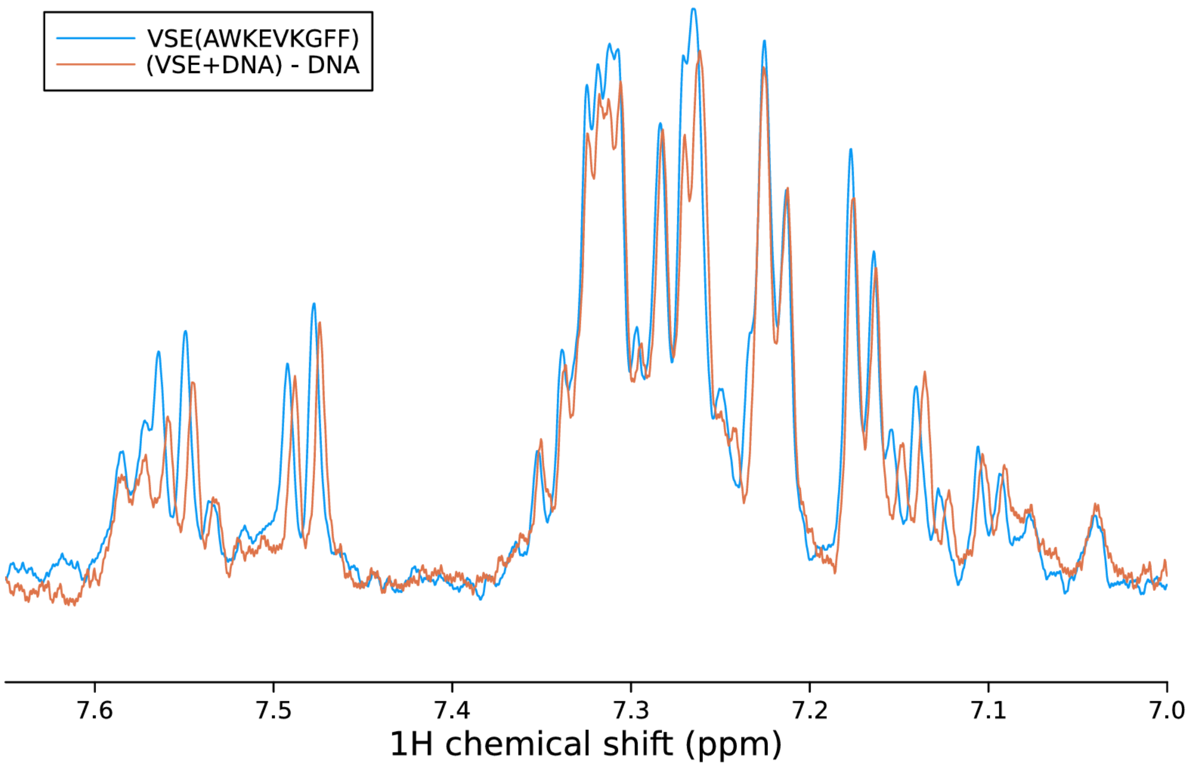

**Figure S20:** $^{1}H$ 1D NMR spectra (600 MHz, 283 K) of Pep-VSE (50 μM, 10 mM NaCaco, pH 6.6, 10% $D_2O$) showing the aromatic region (blue line); a comparison in the presence of 1 equivalent c-MycC52 is shown in red (an equivalent spectrum of DNA alone has been subtracted for clarity).

### 8) Computational Studies

The c-MycC52 iM was selected for molecular dynamics and docking studies. A model of this iM was generated using the crystal structure of the intramolecular iM from the Insulin-Linked Polymorphic Region (ILPR) (PDB ID 8AYG)[1] and the folding pattern proposed by Sutherland *et. al.* based on mutational and bromine footprinting experiments.[2] In this folding pattern the three loops that connect the $CC^+$ core in c-Myc model comprise of 5 nucleotides each (Figure S21). This is distinct from the ILPR structure where each loop is formed from 3 nucleotides. The 5′ and 3′ flanking sequence consists of five and sixteen nucleotides, respectively. The 3′ end flank was trimmed to five nucleotides. The final c-Myc model contains 41 nucleotides with the sequence: 5′- CTTCTCCCCACCTTCCCCACCCTCCCCACCCTCCCCATAAG - 3′

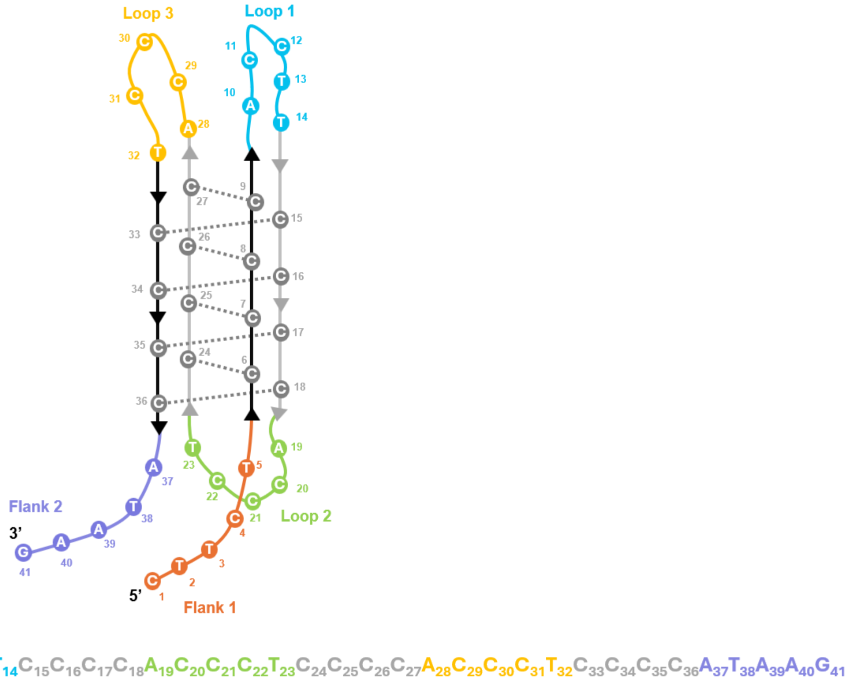

**Figure S21:** Folding pattern used for modelling the c-Myc i-motif coloured based on the flank or loop position in the sequence: flank-1 (orange), loop-1 (cyan), loop-2 (green), loop-3 (yellow), flank-2 (purple), grey (C-core).

The loops in the iM structure have been shown to be dynamic and can adopt multiple conformations.[1] Some conformations have been identified that are potential sites for small molecule and peptide binding.[1] To identify such conformations in the c-Myc model, we carried out enhanced sampling molecular dynamics simulations. Markov state models were subsequently built to cluster metastable states. From these, the most stable states were chosen to carry out molecular docking with the PTN and VSE peptides.

Prior to the simulations, the c-Myc model was prepared using the tleap module in the AmberTools23 package.[3] The AMBER force fields (bsc1 with OL24 modifications) were used to describe the DNA.[4, 5] The parameterisation of iMs requires two of the cytosine stems to be protonated at the N3 position ($C_{6\text{-}9}$ and $C_{15\text{-}18}$). The force field parameters for protonated cytosine were taken from the AMBER force field library (all_prot_nucleic10.lib).

The negatively charged sugar-phosphate backbone was neutralised by adding $K^+$ ions. Additional $K^+$ and $Cl^-$ ions were added to bring the final salt concentration to 0.15 mol/L. The system was solvated using the TIP3P water model in an orthorhombic solvation box, whose edge extended 12 Å from solute. The final dimensions of the solvated box were 68.2 x 59 x 83 $Å^3$.

The system was minimised using 5000 steps of conjugate gradient energy minimisation to relieve any unwanted steric clashes during the preparation of the system. The first step involved the system to be equilibrated under NVT ensemble for 5 ns at 1 atm followed by two consecutive steps of NPT conditions (5 ns each). The temperature was set to 300K. The bonds were set to rigid, a cut off was set to 9 Å and particle mesh Ewalds summation was switched on for long-range electrostatics. During the equilibration, the iM heavy atoms were constrained by a spring constant set at 1 kcal/mol/$Å^2$, which were gradually removed. The ions and the solvent molecules remained free throughout the protocol. The production simulations were run under NVT ensemble using a Langevin thermostat with 1.0 $ps^{-1}$ damping and a hydrogen mass repartitioning scheme to achieve a time step of 4 fs.

The adaptive bandit enhanced sampling protocol was employed to sample the loop and flank conformations.[1, 6] In short, adaptive bandit employs multiple short production runs (50 ns). The objective is to avoid redundant sampling and maximise a reward function. The reward function was defined to be proportional to relative phosphate-phosphate distances of the loops (bases 10-14, 19-23, 28-32). The decision made each time aims to maximise the conformational space explored, where phosphate-phosphate distances are used as a metric. The loops are highly dynamic, able to sample multiple conformations with varying degrees of stability. The system was run for a total of at least 60 μs.

The multiple short trajectories are analysed by generating a Markov State Model (MSM). MSMs allow the integration of multiple simulation trajectories into a single model of the conformational landscape that contains both kinetic and thermodynamic information as well as maintain structural detail. MSMs are built on inter-state transitions; all of these states can be compiled and used to create a single model. Pyemma package is used to build the MSM.[7] The time-lagged independent components analysis (tICA)[8] is used to reduce the dimensionality of the input feature. The alpha and gamma dihedral angles of the loop bases were selected as the input feature. The data from tICA is then projected on the first two ICs at a selected lag time and then clustered using the k-means algorithm.[9] The tICA lag time was chosen to be 80 ns. The number of chosen clusters was 500. This allows the numerous conformations to be mapped onto a set of discrete states and permits MSMs to be generated that describe transitions occurring between any two clusters. The MSM calculates these transitions between states at a given lag time, which was selected to be 1 ns. At this MSM lag time, the underlying processes, which are represented by an implied time scale (ITS) plot, converge (Figure S22A). The MSM is then validated using a Chapman Kolmogorov (CK)[10] test, which tests the ability of the model at longer time scales (Figure S22B). Finally, the Perron cluster-cluster analysis[11] (PCCA++) algorithm is used to coarse grain the conformational space of the metastable states and then approximate the stationary distribution of the data, as well as approximating the relative free energies. A total of 9 discrete states were generated that described different aspects of loop sampling (Figure S23). The most stable conformations (model 8 and model 9) were chosen to carry out molecular docking.

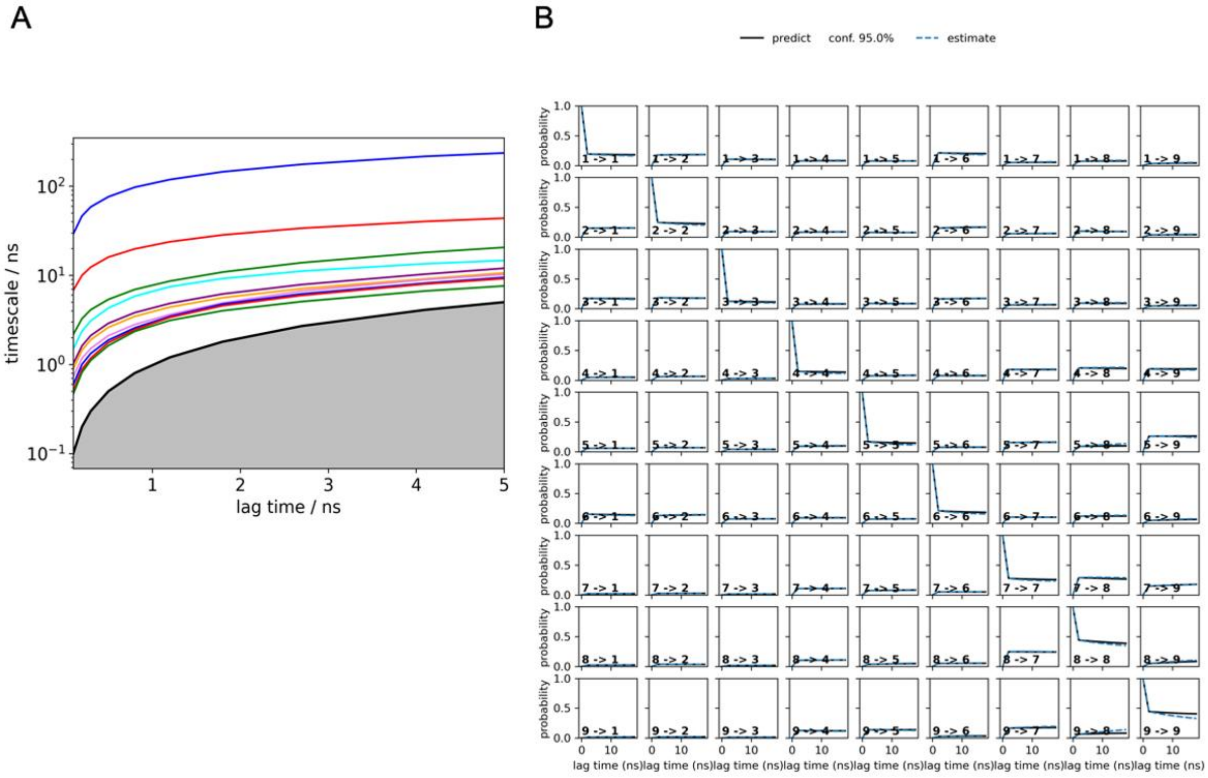

**Figure S22:** (A) The implied timescale (ITS) plot and (B) The Chapman-Kolmogorov (CK) plot is used to validate the built MSM over longer time scales.

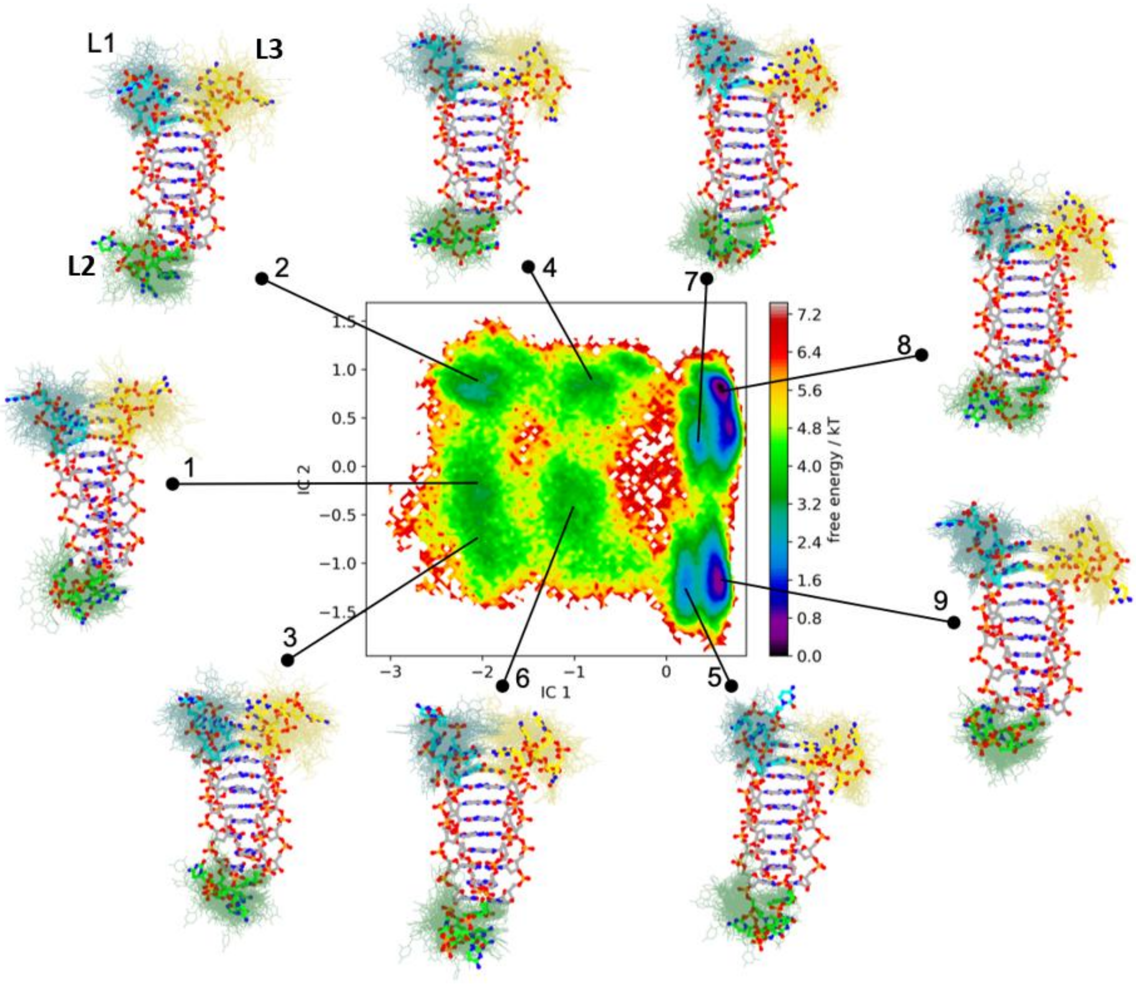

**Figure S23:** Projection of the Markov state model free energy landscape along the time-lagged independent components IC1 and and IC2 in c-Myc i-motif. The represented structures 1-9 derived from the dynamics of the loops are marked in their basins. The positions of the loops have been marked L1 (cyan), L2 (green) and L3 (yellow). The two most stable models 8 and 9 were selected for docking the peptides.

We first performed de novo 3D structure predictions for two peptides, PTN (sequence: PTNVSGRYLFC) and VSE (sequence: VSEAWKEVKGFF), using the PEP-FOLD3 web server (https://bioserv.rpbs.univ-paris-diderot.fr/services/PEP-FOLD3/). Default parameters were utilized, including 100 independent Monte Carlo simulations (30,000 steps each), the fragment-based trajectory (fbt) generator, and the model quality assessment (MQA) option. Generated peptide conformations were evaluated and ranked using the sOPEP coarse-grained force field and subsequently clustered based on backbone RMSD. The centroid structure of the largest cluster for each peptide, representing the most energetically favourable and structurally representative conformation, was selected as the optimal structure for subsequent docking analysis.

Next, molecular docking was performed using the HDOCK web server (http://hdock.phys.hust.edu.cn/) to explore interactions between the peptides and two distinct conformations of the c-Myc DNA (model 8 and model 9). The iM structures in PDB format, were uploaded as receptors, and peptide centroid structures as ligands. Key DNA binding residues (bases from loop 1 and 3) were specified explicitly in the "Binding site residues" field. HDOCK applied its default hybrid docking protocol to generate 100 candidate binding models per docking run, evaluated and ranked using the built-in ITScorePP scoring function and confidence scores. The highest-ranked model (characterised by the most negative docking score and highest confidence score) from each docking experiment was selected as the optimal peptide-DNA complex structure (Figure 3).

Comparative analysis of docking results for i-motif model 8 and model 9 with peptides Pep-PTN and Pep-VSE revealed notable differences in binding affinities. Model 9 consistently yielded superior docking scores and confidence scores for both PTN (–159.94, 0.55) and Pep-VSE (–154.33, 0.52) compared to Model 8 (Pep-PTN: –152.09, 0.51; Pep-VSE: –147.54, 0.49). Additionally, Pep-PTN exhibited slightly stronger binding interactions compared to Pep-VSE across both DNA conformations. These results indicate that model 9 forms more stable peptide-i-motif complexes, with Pep-PTN demonstrating marginally enhanced binding capability.
